# Feature-specific salience maps in human cortex

**DOI:** 10.1101/2023.03.29.534828

**Authors:** Daniel D. Thayer, Thomas C. Sprague

## Abstract

Priority maps are representations of visual space that determine the relative importance of scene locations. Computational theories suggest that priority maps identify salient locations based on individual feature dimensions (e.g., color, motion), which are integrated into an aggregate priority map. While widely accepted, a core assumption of this framework—the existence of independent feature dimension maps in visual cortex—remains untested. Here, we tested the hypothesis that feature-selective retinotopic regions in human cortex act as neural feature dimension maps, indexing salience based on their preferred feature. We used fMRI activation patterns to reconstruct spatial maps while participants viewed stimuli with salient regions defined by color or motion direction. Reconstructed spatial maps selectively represented salient locations defined by each region’s preferred feature. These findings identify spatially organized feature dimension maps that characterize the salience of scene locations based on a specific visual feature, confirming a key prediction of priority map theory.

## Introduction

Often, we must search for items that are relevant to ongoing goals, such as the coffee maker in the morning or the box of cookies after dinner. However, objects within a given scene are constantly vying for our attention. A salient but task-irrelevant object, like a bright yellow banana on the counter or a cat leaping across the kitchen, may distract our attention and slow the search for coffee (or a cookie). One prominent model that highlights the competition between task-irrelevant salience and task-relevant goals in guiding attention is priority map theory (Awh et al., 2012; Fecteau & Munoz, 2006; Itti & Koch, 2001; Serences & Yantis, 2006; Treisman & Gelade, 1980; Wolfe, 1994). Per this theory, the visual system computes a priority map, which is a representation of visual space indexing the relative importance—or *priority—*of locations in the environment. Priority is computed based on both salience—defined based on image-computable properties—and relevance—defined by an individual’s current goals, and is used to direct attention to the highest-priority locations for further processing (Carrasco, 2011; Eckstein, 2011; Yu et al., 2023).

To compute the salience associated with a given location, priority map theory posits that information about individual feature dimensions (e.g., color, motion, etc) is independently extracted from retinal input into a series of ‘feature dimension maps’. For a given feature dimension map, salient regions of space are defined based on within-dimension local feature contrast, such as the aberrant color of the yellow banana or motion direction of the leaping cat (Itti & Koch, 2000, 2001). A conspicuous location defined by a given feature dimension is given high activation in the corresponding feature dimension map (the yellow banana would result in strong activation in a color map, but not a motion map; Fig. 1). Activity profiles across various feature dimension maps are then integrated into a unified feature-agnostic priority map, which indexes the most important locations within the visual field, regardless of the source of their importance.

**Figure 1.**
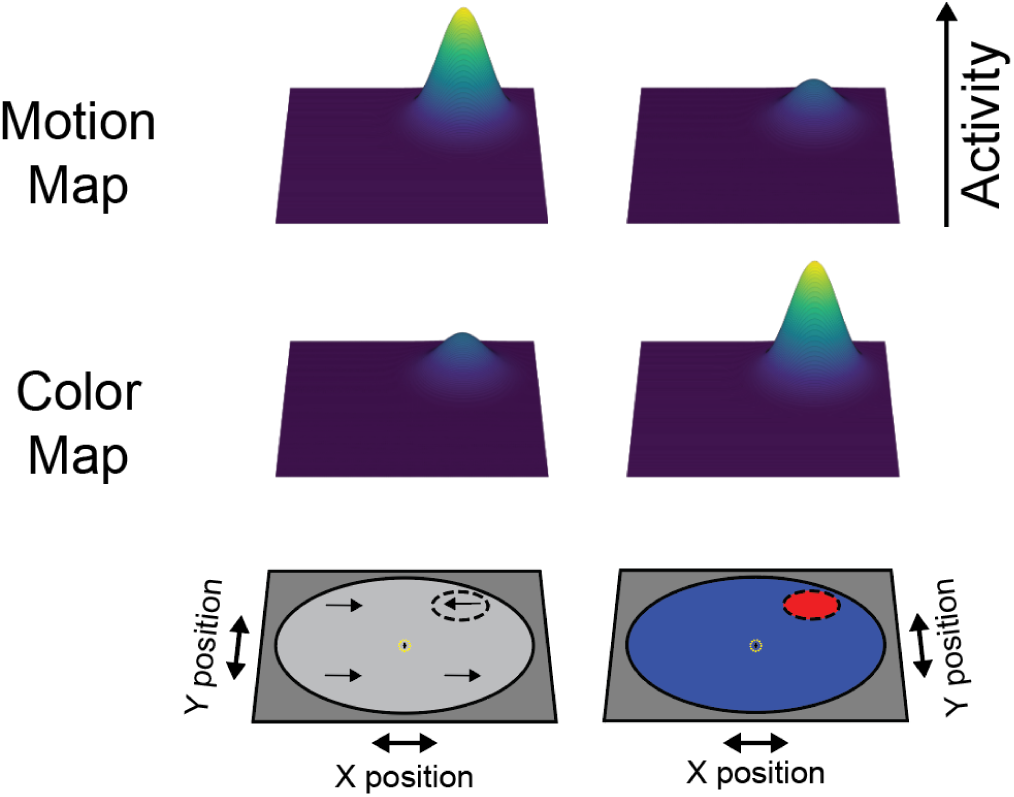
Feature dimension maps index salient locations based on their preferred feature dimension. Priority map theory invokes ‘feature dimension maps’ to compute salient location(s) based on local feature contrast within each feature dimension (e.g., color, motion). Accordingly, when a location in a stimulus display is made salient based on local differences in motion direction, activation profiles over a ‘motion map’ should track the salient location, while a ‘color map’ would not. Similarly, map activation corresponding to a salient motion stimulus should be stronger in a ‘motion map’ than when a location is made salient based on local differences in color. Complementary results would be predcted for a ‘color map’, with stronger activation at the location of a salient color stimulus as compared to a salient motion stimulus. While these feature dimension maps productively account for behavioral results in visual search tasks, it remains unknown whether salient locations are independently indexed in different feature-selective regions in visual cortex.

Studies in humans and nonhuman primates have identified stronger neural responses associated with salient stimulus locations than those associated with non-salient locations in visual (Bogler et al., 2013; Burrows & Moore, 2009; Mulckhuyse et al., 2011; O’Connell & Chun, 2018; Ogawa & Komatsu, 2004, 2006; Poltoratski et al., 2017; Poltoratski & Tong, 2020; Yildirim et al., 2018), parietal (Bisley & Goldberg, 2006; Bogler et al., 2011; Chen et al., 2020; Gottlieb et al., 1998), and frontal cortex (Bichot & Schall, 1999; Schall et al., 1995; Schall & Hanes, 1993), and in subcortical structures including the superior colliculus (White et al., 2017), consistent with a neural instantiation of a priority map (Bisley & Goldberg, 2010; Fecteau & Munoz, 2006; Katsuki & Constantinidis, 2014; Serences & Yantis, 2006). However, despite this converging neural evidence offering strong support for the implementation of feature-agnostic priority maps across the brain, support for feature dimension maps is primarily based on behavioral studies measuring visual search response times (Bacon & Egeth, 1994; Huang & Pashler, 2007; Müller et al., 1995; Theeuwes, 1992; Treisman, 1998; Wolfe & Horowitz, 2004). Indeed, neural studies have all either focused on stimulus displays involving a single salient feature dimension (Beck & Kastner, 2005; Cook & Maunsell, 2002; Moran & Desimone, 1985; Zhang et al., 2012), measured from a single feature-selective visual region (Burrows & Moore, 2009; Martıń ez-Trujillo & Treue, 2002; Mazer & Gallant, 2003; Ogawa & Komatsu, 2004, 2006; Reynolds & Desimone, 2003), or studied feature-agnostic salience (Bisley & Goldberg, 2006; Bogler et al., 2011, 2013; Gottlieb et al., 1998; Sprague, Itthipuripat, et al., 2018). Thus, despite the key theoretical role specific stimulus feature dimensions are believed to play in computing representations of stimulus salience, it remains unknown whether the brain implements this compartmentalized computational architecture.

Here, we sought to resolve this question by testing the hypothesis that feature-selective retinotopic regions of visual cortex preferentially index salient locations based on their preferred feature dimension. We focused on regions exhibiting both retinotopic organization defined using an independent retinotopic mapping task and feature selectivity verified with independent functional localizers. Within these candidate ROIs, we characterized feature selectivity and spatial salience computations for task-irrelevant visual stimuli defined by different feature dimensions using a multivariate image reconstruction technique. Participants attended a central fixation point while viewing stimulus displays typically containing one salient stimulus location. Across trials, we varied the salience-defining stimulus feature (color, motion, or a single salient stimulus in isolation). If color-selective retinotopic regions hV4/VO1/VO2 (Brewer et al., 2005; Conway et al., 2007; Mullen, 2019) and motion-selective retinotopic regions TO1/TO2 (Albright, 1993; Amano et al., 2009; Huk et al., 2002) act as neural feature dimension maps, salient stimuli will result in patterns of multivariate BOLD responses containing strong activation at the salient location, and these representations will be strongest when the salient location is defined by a region’s preferred feature dimension (Fig. 1). Alternatively, if extrastriate cortex does not compartmentalize salience computations based on individual feature dimensions, retinotopic ROIs should equivalently index salience regardless of its defining feature dimension. Consistent with the predictions of priority map theory, we observed strong representations of stimuli localized to the salient stimulus location in the display. Moreover, the strength of the stimulus representation was strongest in the ROI selective for the salience-defining feature. Together, these results suggest that these extrastriate visual regions serve to extract salient locations based on local feature contrast within their preferred feature dimensions, supporting their role as neural feature dimension maps.

## Materials & Methods

### Participants

8 subjects recruited from the University of California, Santa Barbara (UCSB) community participated in the study (6 female, 18-27 years old). Pilot data (n = 3) confirmed that this sample size allowed for adequate power to detect our effects of interest (d_z_ = 3.10). We opted to collect a large number of measurements from each subject to minimize within-subject variance, which often benefits statistical power more than increased sample sizes (Baker et al., 2021). All subjects reported normal or corrected-to-normal vision and did not report neurological conditions. Procedures were approved by the UCSB Institutional Review Board (IRB# 2-20-0012). All subjects gave written informed consent before participating and were compensated for their time ($20/h for scanning sessions, $10/h for behavioral familiarization/training).

### Stimuli and Procedure

Participants performed a 30-minute training session before scanning so that they were familiarized with the instructions. We used this session to establish the initial behavioral performance thresholds used in the first run of the scanning session. In the main task session, we scanned participants for a single two-hour period consisting of at least 4 mapping task runs, which we used to independently estimate encoding models for each voxel, and 8 experimental feature-salience task runs. All participants also underwent additional anatomical and retinotopic mapping scanning sessions (1-2x 1.5-2 hr sessions) to identify regions of interest (ROI; see Region of interest definition). Additionally, most participants (n = 6) underwent an independent functional localizer session which we used to verify retinotopically defined ROIs were feature selective.

Stimuli were presented using the Psychophysics toolbox (Brainard, 1997; Pelli, 1997) for MATLAB (The MathWorks, Natick, MA). Visual stimuli were rear-projected into a screen placed ∼110 cm from the participant’s eyes at the head of the scanner bore using a contrast-linearized LCD projector (1,920×1,080, 60 Hz) during the scanning session. In the behavioral familiarization session, we presented stimuli on a contrast-linearized LCD monitor (2,560×1,440, 60 Hz) 62 cm from participants, who were seated in a dimmed room and positioned using a chin rest. For all sessions and tasks (main tasks, localizers, and mapping task), we presented stimuli on a neutral gray circular aperture (9.5° radius), surrounded by black (only aperture shown in Fig. 1).

### Feature-salience task

For the main task (Fig. 2) and functional localizers (Fig. S1; see below), participants attended a flashing cross within the fixation circle and ignored any other stimuli presented throughout the scanning session. This task localized goal-directed attention to fixation and was equivalent across all stimulus conditions, allowing us to isolate signals associated with bottom-up salience processing of our peripheral stimuli. Participants monitored the fixation cross throughout the whole run for any increase in length in either the vertical or horizontal bar of the cross and responded to changes with a button press (the left button for a horizontal target, the right button for a vertical target). The vertical and horizontal lines of the fixation cross were 0.25° of visual angle long and flickered at 3 Hz (10 frames on, 10 frames off at 60 Hz). Whenever the cross was visible, there was a 22.5% chance either line had a small change in length. When a change was detected, participants reported which line increased in length (horizontal or vertical). To ensure participants maintained vigilant attention at fixation throughout the entire experiment, we adjusted the difficulty of the fixation task between runs by altering the degree of size change for vertical/horizontal lines based on behavioral accuracy (range: 0.05° to 0.125°). Participants performed the fixation task continuously throughout both stimulus presentation periods and ITIs to ensure salient events were temporally decoupled from fixation task performance and/or target detection. Feedback for each response to the fixation task was given via the aperture around fixation changing color, with green indicating a correct response, red indicating an incorrect response, and yellow indicating no response. A fixation target was never present for the first or last 2 s of a trial, or for 2 s after the presentation of a previous target.

**Figure 2.**
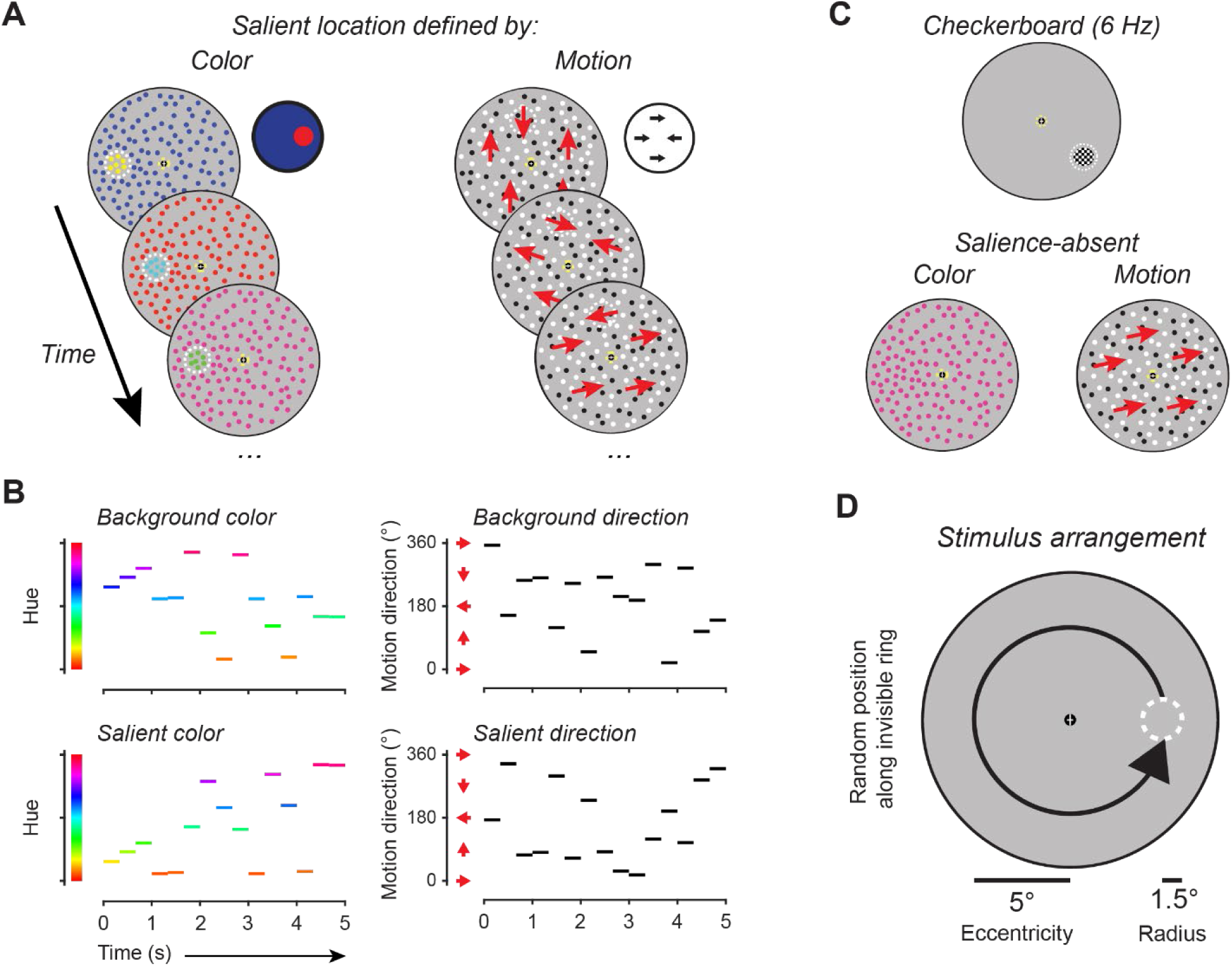
Feature-salience task. **A:** On each fMRI scanning run, participants continuously performed an attention-demanding task at fixation where they reported changes in length of either the horizontal or vertical bar of the fixation cross. While attention was directed to the demanding fixation task, we measured how feature-selective retinotopic ROIs encode task-irrelevant salient stimulus locations by presenting various types of visual stimuli. On most stimulus presentation trials, the visual stimulus consisted of dots spanning the entire screen. The dot stimuli could either be presented as static colored dots, or grayscale (black/white) moving dots. **B:** The features of all dots were updated at 3 Hz such that, on average, the overall feature value (color/motion) presented across each trial was neutral. For example, on ‘color’ trials, the color and location of each static dot was updated every 333 ms with a new randomly-selected hue and randomly-drawn location. On most dot array trials (66.6%; ‘salience-present’), a circular portion of the stimulus display was made salient by presenting dots in the opposite feature value as presented in the background. For example, during a 333 ms period, if the background dots were moving at 45°, the salient foreground dots would be moving 225°. These salient stimulus regions were never relevant for participant behavior, and the challenging fixation task ensured attention was withdrawn from peripheral stimuli. After stimulus presentation, there was a 6-9 s blank ITI during which time the fixation task continued. Example trial for each condition shown. **C:** As control conditions, we also included trials with the salient location defined by a flickering checkerboard (6 Hz full-field flicker) on a blank background, and trials with colored static or moving black and white dots with no salient location. **D:** On salience-present trials, the salient stimulus was 1.5° in radius, and was presented at a location randomly chosen from an invisible ring centered 5° from fixation.

The critical, ignored, stimuli were either a color- or motion-defined salient location presented as a circular disc within random dot arrays spanning the entire stimulus aperture except for a region around fixation (0.75°). On color trials, static dots within a disc were presented in the ‘opposite’ color (HSV colorspace) as compared to the background dots; on motion trials, moving black and white dots within a disc were presented in the opposite motion direction as compared to the background dots. For example, if the dot array contained dots moving at 0° (to the right), the motion-defined salient location would contain dots moving at 180° (to the left). Individual dots occupied 0.05° of visual angle, and dot density was 15 dots/deg^2^. In the motion array, dots moved at a speed of 9.5° /s in a randomly selected planar direction and each dot was randomly colored black or white (100% contrast). Dots were randomly replotted every 50 ms or when they exceeded the stimulus bounds. For the color array, all dots remained static and were assigned a random hue value. Dot locations were updated every 333 ms. Both arrays updated every 333 ms during the 5 s presentation period, such that a new color or motion value was applied to every dot in the updated array three times per second. Trials started with the onset of the peripheral dot array while participants were attending fixation. The salient location appeared throughout the entire stimulus interval, centered 5° from fixation at a random location along an invisible ring from 0°-359° and had a radius of 1.5°.

We included 3 additional control conditions intermixed with salience-present trials. First, to ensure spatially localized activation is due to the presence of the salient location, we presented colored static dots and moving black and white dots with no salient location defined (‘salience-absent’ trials). Second, as a positive control to ensure our image reconstruction procedure was effective in each retinotopic ROI, we presented a flickering checkerboard disc (spatial frequency 0.679 cycles/°) on a gray background at the same size, eccentricity, and duration as the salient discs (‘checkerboard’ trials; similar to previous reports; Sprague & Serences, 2013). The checkerboard stimulus flickered at a rate of 6 Hz and was considered to be feature-agnostic with respect to the key manipulations in the study (i.e., color/motion). All trials were separated by a randomly selected ITI ranging from 6-9 s with an average ITI of 7.5 s.

Each run had 24 trials. There were 6 trials of each salience-present condition (based on color, based on motion, checkerboard) and 3 trials each of the salience-absent color/motion conditions. Trial order was shuffled within run. Each run started with a 3 s blank period and ended with a 10.5 s blank period, for a total run duration of 313.5 s (for one participant, we acquired 416 TRs for one run instead of 418, resulting in 312 s of data for this run). Eye position was monitored throughout the experiment using an Eyelink 1000 eyetracker (SR Research).

### Spatial mapping task

We also acquired several runs of a spatial mapping task used to independently estimate a spatial encoding model for each voxel, following previous studies (Sprague et al., 2016; Sprague, Itthipuripat, et al., 2018; Sprague & Serences, 2013). On each trial of the mapping task, we presented a flickering checkerboard at different positions selected from a hexagonal grid spanning the screen. Participants viewed these stimuli and responded whenever a rare contrast change occurred (10 out 47 trials, 21.3%), evenly split between contrast increments and decrements. The checkerboard stimulus was the same size as the salient locations in the feature salience task (1.5° radius) and was presented at 70% contrast and 6-Hz full-field flicker. All stimuli appeared within a gray circular aperture with a 9.5° radius, as in the feature-salience task. For each trial, the location of the stimulus was selected from a triangular grid of 37 possible locations with an added random uniform circular jitter (0.5° radius). The base position of the triangular grid was rotated by 30° on every other scanner run to increase spatial sampling density. As a result, every mapping trial was unique, which enabled robust spatial encoding model estimation.

Each trial started with a 3000 ms stimulus presentation period. If a target was present, then the stimulus would be dimmed/brightened for 500 ms with the stipulation that the contrast change would not occur in either the first or last 500 ms of the stimulus presentation period. Finally, there was an ITI ranging from 2 to 6 s (uniformly sampled using the linspace command in MATLAB [linspace(2, 6, 47)]). All target-present trials were discarded when estimating the spatial encoding model. Each run consisted of 47 trials (10 of which included targets). We also included a 3 s blank period at the beginning of the run and a 10.5-s blank period at the end of the run. Each run totaled 432 s.

#### Retinotopic mapping task

We used a previously reported task (Mackey et al., 2017) to identify retinotopic regions of interest (ROIs) via the voxel receptive field (vRF) method (Dumoulin & Wandell, 2008). Each run of the retinotopy task required participants attend several random dot kinematograms (RDK) within bars that would sweep across the visual field in 2.25 s (or, for one participant, 2.6 s) steps. Three equally sized bars were presented on each step and the participants had to determine which of the two peripheral bars the motion in the central bar matched with a button press. Participants received feedback via a red or green color change at fixation. We used a three-down/one-up staircase to maintain ∼80% accuracy throughout each run so that participants would continue to attend the RDK bars. RDK bars swept 17.5° of the visual field. Bar width and sweep direction was pseudo-randomly from several different widths (ranging from 2.0° to 7.5°) and four directions (left-to-right, right-to-left, bottom-to-top, and top-to-bottom).

### Functional localizer tasks

To independently identify color- and motion-selective voxels, we scanned participants while they performed 3 runs each of a color and motion localizer task (Bartels & Zeki, 2000; Huk et al., 2002) using a blocked design. During both tasks, the participant performed the same fixation task from the feature-salience attention task, where they monitored a central cross for changes in the size of the horizontal and vertical lines. In the color localizer task, participants viewed colored or greyscale rectangles of various sizes within the same aperture dimensions described in the spatial mapping task (see spatial mapping task). Stimuli were presented spanning the entire aperture. Rectangle colors were individually sampled from the entire RGB colorspace (uniform independent random distribution of R, G, and B). Similarly, each greyscale rectangle had a randomly sampled contrast (identical R, G, and B value, randomly sampled for each rectangle). During the motion localizer, participants viewed either static or moving black and white dots. For the moving dots, motion could be CW, CCW, or planar (20 evenly spaced steps from 18 to 360°). Dots were redrawn every 100 ms or when they exceeded the stimulus boundary. When the array contained planar motion, dots moved at 1.2°/s; if CW/CCW motion, 0.6°/s. Within each block, each stimulus (static/motion or color/greyscale) was shown for 400 ms followed by a 100 ms blank period before the next stimulus presentation. Each block lasted 18 s (36 updates of stimulus feature values), with feature values randomly selected each presentation. During each scanning run, we presented 6 total blocks, alternating between grayscale rectangles (static dots) and colored rectangles (moving dots) for the color (motion) localizer runs. At the end of each run, participants viewed a blank screen while performing the fixation task for 18 s. Runs started with a 3 s blank period and ended with a 10.5 s blank period. There was no fixation task during the start and end blank periods. Each run lasted 229.5 s.

We acquired localizer data for 6 of 8 participants (the other 2 participants were unable to return to complete the localizer session).

### fMRI acquisition

fMRI data acquisition and preprocessing pipelines in the current study closely followed a previous report (Hallenbeck et al., 2021) but with slight modifications. We acquired all functional and anatomical images at the UCSB Brain Imaging Center using a 3T Siemens Prisma scanner. fMRI scans for experimental, model estimation, retinotopic mapping, and functional localizers were acquired using the CMRR MultiBand Accelerated EPI pulse sequences. We acquired all images with the Siemens 64 channel head/neck coil with all elements enabled. We acquired both T1- and T2- weighted anatomical scans using the Siemens product MPRAGE and Turbo Spin-Echo sequences (both 3D) with 0.8 mm isotropic voxels, 256 x 240 mm slice FOV, and TE/TR of 2.24/2400 ms (T1w) and 564/3200 ms (T2w). We collected 192 and 224 slices for the T1w and T2w, respectively. We acquired three T1 images, which were aligned and averaged to improve signal-to-noise ratio.

For all functional scans, we used a Multiband (MB) 2D GE-EPI scanning sequence with MB factor of 4, acquiring 44 2.5 mm interleaved slices with no gap, isotropic voxel size 2.5 mm and TE/TR: 30/750ms, and P-to-A phase encoded direction to measure BOLD contrast images. For retinotopic mapping of one participant (sub004), we used a MB 2D GE-EPI scanning sequence acquired 56 2 mm interleaved slices with isotropic voxel size 2 mm and TE/TR: 42/1300 ms. We measured field inhomogeneities by acquiring spin echo images with normal and reversed phase encoding (3 volumes each), using a 2D SE-EPI with readout matching that of the GE-EPI and the same number of slices, no slice acceleration, TE/TR: 45.6/3537ms (TE/TR: 71.8/6690ms for sub004’s retinotopic mapping session).

### MRI preprocessing

Our approach for preprocessing was to coregister all functional images to each participant’s native anatomical space. First, we used all intensity-normalized high-resolution anatomical scans (3 T1 images and 1 T2 image for each participant) as input to the ‘hi-res’ mode of Freesurfer’s recon-all script (version 6.0) to identify pial and white matter surfaces. Processed anatomical data for each participant was used as the alignment target for all functional datasets which were kept within each participant’s native space. We used AFNI’s afni_proc.py to preprocess functional images, including motion correction (6-parameter affine transform), unwarping (using the forward/reverse phase-encode spin echo images), and coregistration (using the unwarped spin-echo images to compute alignment parameters to the anatomical target images). We projected data to the cortical surface, then back into volume space, which incurs a modest amount of smoothing perpendicular to the cortical surface. To optimize distortion correction, we divided functional sessions into 3-5 sub-sessions which consisted of 1-4 fMRI runs and a pair of forward/reverse phase encode direction spin echo images each, which were used to compute that sub-session’s distortion correction field. For the feature salience and mapping task, we did not perform any spatial smoothing beyond the smoothing introduced by resampling during coregistration and motion correction. For retinotopic mapping and functional localizer scans, we smoothed data by 5 mm FWHM on the surface before projecting back into native volume space.

### Region of interest definition

We identified 15 ROIs using independent retinotopic mapping data. We fit a vRF model for each voxel in the cortical surface (in volume space) using averaged and spatially smoothed (on the cortical surface; 5 mm FWHM) time series data across all retinotopy runs (8-12 per participant). We used a compressive spatial summation isotropic Gaussian model (Kay, Winawer, Mezer, et al., 2013; Mackey et al., 2017) as implemented in a customized, GPU-optimized version of mrVista (see Mackey et al., 2017) for detailed description of the model). High-resolution stimulus masks were created (270 x 270 pixels) to ensure similar predicted responses within each bar size across all visual field positions. Model fitting began with an initial high-density grid search, followed by subsequent nonlinear optimization. We visualized retinotopic maps by projecting vRF best-fit polar angle and eccentricity parameters with variance explained ≥10% onto each participant’s inflated cortical surfaces via AFNI and SUMA (Fig. 3; Fig. S2). We drew retinotopic ROIs (V1, V2, V3, V3AB, hV4, LO1, LO2, VO1, VO2, TO1, TO2, IPS0-3) on each hemisphere’s cortical surface based on previously-established polar angle reversal and foveal representation criteria (Amano et al., 2009; Mackey et al., 2017; Swisher et al., 2007; Wandell et al., 2007; Winawer & Witthoft, 2015). Finally, ROIs were projected back into volume space to select voxels for analysis.

**Figure 3.**
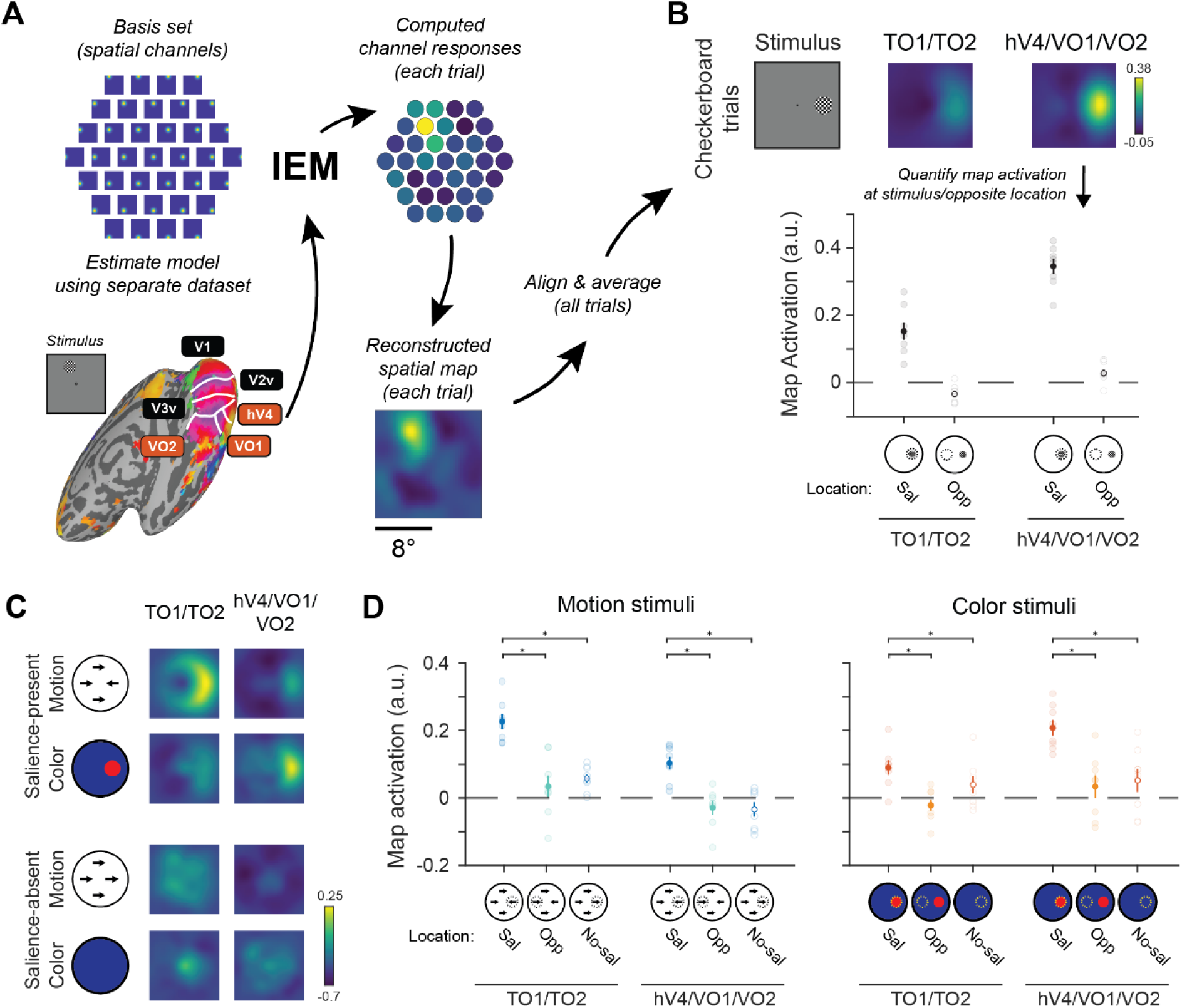
Reconstructed spatial maps track salient stimulus location. **A:** We estimated a spatial inverted encoding model (IEM) for each ROI using an independent spatial mapping task (see Methods for details). Using this spatial encoding model, which maps activation patterns to activation of spatial channels that can be summed to produce reconstructed spatial maps, we were able to generate image reconstructions of the visual field on each trial and directly compare map activation across conditions. For each condition, we averaged trial-wise reconstructions computed using activation patterns from 5-8 s after stimulus onset after we rotated and aligned them to the known position of the salient stimulus, if present. **B.** To validate the utility of our method, we computed reconstructions of checkerboard trials from each ROI. Qualitatively, there was a strong response to the checkerboard stimulus across both aggregate ROIs. To quantify reconstructions, we computed the mean map activation at the aligned stimulus location and at the location on the opposite side of fixation within each ROI. In each ROI’s reconstruction, activation at the stimulus location was greater than at the opposite location. two-way permuted repeated-measures ANOVA (ROI; location) identified a significant main effect of location (*p* < 0.001). **C.** Qualitatively, the salient location was highlighted in the aggregate motion ROI TO1/TO2 when the salient location was defined by motion, but not by color, with the converse result observed in the aggregate color ROI hV4/VO1/VO2. On salience-absent trials, no location is reliably highlighted in average reconstructions. **D:** Using data from each stimulus condition, we identified whether enhanced reconstruction responses were localized to the salient stimulus by comparing mean salient location activation (‘Sal’) to the mean activation of the position opposite of the salient location (‘Opp’), as well as the mean activation of the ‘aligned’ position of the salience-absent condition (‘No-sal’). On trials with motion-defined stimuli, activation in the motion-selective ROI was greatest at the location of the salient motion stimulus. When stimuli were defined by static colorful dots, activation in the color-selective ROI was greatest at the location of the salient color stimulus. In all panels, error bars are SEM across participants (n = 8). * indicates signififant difference based on permuted paired T-test, corrected for multiple comparisons with FDR. three-way permuted repeated-measures ANOVA (ROI; location; stimulus feature) identified a significant main effect of location (p < 0.001), two-way interaction between feature and ROI (p = 0.007), and a three-way interaction (p = 0.001). For individual ROI results, see Figure S3. For all statistical comparisons, see Table S3.

For primary analyses (Figs 3-4), we aggregated across color-selective maps (hV4, VO1, VO2) and motion-selective maps (TO1, TO2) by concatenating all voxels for which the best-fit vRF model explained at least 10% of the signal variance prior to univariate or multivariate analyses. We also conducted all analyses for each individual ROI using the same voxel selection threshold (see Supplemental Information).

**Figure 4.**
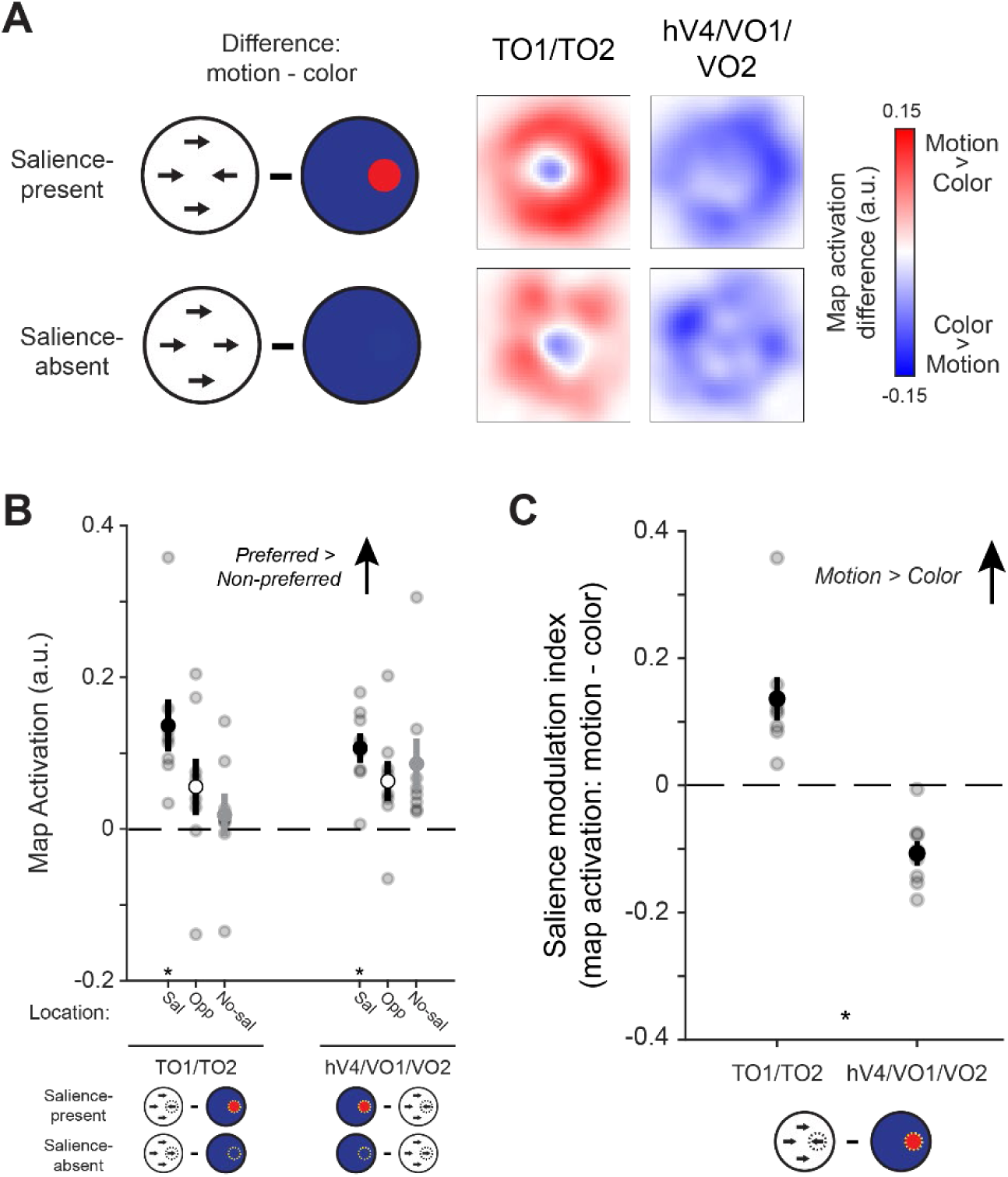
Neural dimension maps selectively index salience based on their preferred feature. **A:** We directly compared reconstructed spatial maps for each ROI between trials where the salient location was defined by motion to those where the salient location was defined by color by computing their pixelwise difference (motion – color). For comparison, we also computed these maps for salience-absent trials. Positive values indicate a region more stronlgy represents a location based on motion stimuli over color stimuli, and negative values indicate the opposite. **B:** To compare feature selectivity across spatial locations and salience-presence conditions, we extracted values of each ROI’s difference map computed between the preferred and non-preferred feature dimension for that ROI (see cartoons). Difference map activation was more positive (preferred > non-preferred) at the salient location than the opposite location on salience-present trials, and than a random location on salience-absent trials (two-way permuted repeated-measures ANOVA with factors of location and ROI; significant main effect of location *p* = 0.002). Additionally, difference map values were only reliably greater than zero at the aligned position on salience-present trials (one-sample T-tests against zero, FDR corrected, *p* ≤ 0.005), indicating that these ROIs preferentially encode salient locations based on their preferred feature dimension. * indicates signifiant difference from zero. **C:** Difference map activation (A) computed at the salient location reliably differed between ROIs, such that motion-selective TO1/TO2 indexed salience more strongly when it was defined by motion than by color, and vice versa for color-selective hV4/VO1/VO2. Asterisk indicates significant difference based on permuted paired-samples T-test, *p* < 0.001. Error bars reflect SEM across participants.

### Inverted encoding model

We used a spatial inverted encoding model (IEM) to reconstruct images based on stimulus-related activation patterns measured across entire ROIs (Sprague & Serences, 2013; Fig. 2A). To do this, we first estimated an encoding model, which describes the sensitivity profile over the relevant feature dimension for each voxel in a region. This requires using data set aside for this purpose, referred to as the “training set”. Here, we used data from the spatial mapping task as the independent training set. The encoding model across all voxels within a given region is then inverted to estimate a mapping used to transform novel activation patterns from a “test set” (runs from the feature salience task) and reconstruct the spatial representation of the stimulus at each timepoint.

We built an encoding model for spatial position based on a linear combination of 37 spatial filters (Sprague et al., 2014; Sprague, Itthipuripat, et al., 2018; Sprague & Serences, 2013). Each voxel’s response was modeled as a weighted sum of each identically shaped spatial filter arrayed in a triangular gird (Fig. 2A). The centers of each filter were spaced by 2.83° and were Cosine functions raised to the 7^th^ power:

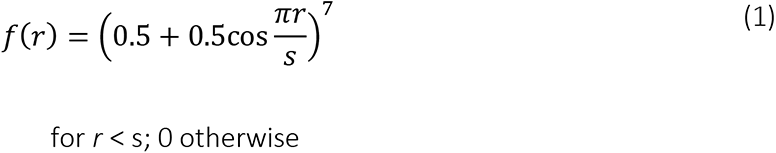

where *r* is the distance from the filter center and *s* is a size constant. The size constant reflects the distance from the center of each spatial filter at which the filter returns to 0. This triangular grid of filters forms the basis set, or information channels for our analysis. For each stimulus used in our mapping task, we converted from a contrast mask to a set of filter activation levels by taking the dot product of the vectorized stimulus mask (*n* trials × *p* pixels) and the sensitivity profile of each filter (*p* pixels × *k* channels). We then normalized the estimated filter activation levels such that the maximum activation was 1 and used this output as *C_1_* in the following equation, which acts as the forward model of our measured fMRI signals:

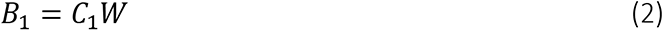

***B_1_*** (*n* trials × *m* voxels) in this equation is the measured fMRI activity of each voxel during the visuospatial mapping task and ***W*** is a weight matrix (*k* channels × *m* voxels) which quantifies the contribution of each information channel to each voxel. ***Ŵ*** can be estimated using ordinary least-squares linear regression to find the weights that minimize the differences between predicted values of ***B*** and the observed ***B_1_***:

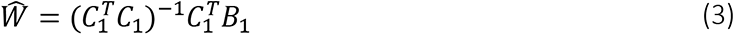

This is computed for each voxel within a region independently, making this step univariate. The resulting *Ŵ* represents all estimated voxel sensitivity profiles. We then used *Ŵ* and the measured fMRI activity of each voxel (i.e., BOLD response) during each trial (using each TR from each trial, in turn) of the feature salience task using the following equation:

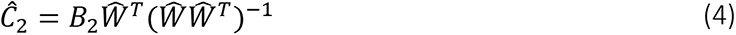

Here, ***Ĉ***_**2**_ represents the estimated activation of each information channel (*n* trials × *k* channels) which gave rise to that observed activation pattern across all voxels within a given ROI (***B_2_***; *n* trials × *m* voxels). To aid with visualization, quantification, and coregistration of trials across stimulus positions, we computed spatial reconstructions using the output of Equation 4. To do this, we weighted each filter’s spatial profile by the corresponding channel’s reconstructed activation level and then summed all weighted filters together (Fig. 3A).

Since stimuli in the feature-selective attention task were randomly positioned on every trial, we rotated the center position of spatial filters such that the resulting 2D reconstructions of the stimuli were aligned across trials and participants (Fig. 3B). We then sorted trials based on condition (salience-present: color, salience-present: motion, checkerboard, salience-absent: color, salience-absent: motion. Finally, we averaged the 2D reconstructions across trials within the same condition for individual participants, then across all participants for our grand-average spatial reconstructions (Fig. 3B-C; Fig. S3). Individual values within the 2D reconstructed spatial maps correspond to visual field coordinates. To visualize feature-selectivity within reconstructed spatial maps, we computed the difference in map activation between the salience-present: color and motion conditions (Fig. 4A). We used these difference maps to assess whether feature-selective ROIs had the same feature preferences throughout the visual field, or if they were localized to the position of the salient stimulus when present.

Critically, because we reconstructed all trials from all conditions of the feature-selective attention task using an identical spatial encoding model estimated with an independent spatial mapping task, we can compare reconstructions across conditions on the same footing (Sprague, Adam, et al., 2018; Sprague et al., 2019; Sprague, Itthipuripat, et al., 2018). Moreover, because we were not interested in decoding precision, but instead in the activation profile across the entire reconstructed map, we did not employ any feature decoding approaches and instead opted to directly characterize the resulting model-based reconstructions (e.g., correlation table; Scotti et al., 2021). Finally, the resulting model-based reconstructions are necessarily based on the modeling choices used here and should not be used to infer any features of single-neuron tuning properties (which we do not claim in this report; Sprague, Adam, et al., 2018; Sprague et al., 2019). Should readers be interested in testing the impact these modeling choices have on results, all analysis code and data [will be] freely available (see below).

### Quantifying stimulus representations

To quantify the strength of stimulus representations within each reconstruction, we computed the mean map activation of pixels located within a 1.5° radius disk centered at the known position of each stimulus (matching the stimulus radius of 1.5°; see Sprague, Itthipuripat, et al., 2018). This provides a single value corresponding to the activation of the salient stimulus location for a given condition, within each retinotopic ROI. To assess the spatial selectivity of reconstructed spatial maps, we compared the mean map activation at the location of salient stimuli to map activation at the location opposite fixation using a 1.5° radius disk (Fig. 3B, 3D; Fig. S3).

To compare values across conditions, we computed a difference in extracted map activation across conditions (e.g., subtracted color map activation from motion map activation, Fig. 4). As an exploratory analysis, we computed these differences at every pixel in reconstructed maps. For quantification, we focused on activation averaged over the discs aligned to the salient stimulus location within each map or the opposite location (Fig. 4C).

### Statistical analysis

We used parametric statistical tests for all comparisons (repeated-measures ANOVAs and T-tests). To account for possible non-normalities in our data, we generated null distributions for each test using a permutation procedure (see below) to derive p-values.

First, we used a one-way repeated-measures ANOVA (factor: stimulus condition; 5 levels: motion salience, color salience, checkerboard, non-salient motion, and non-salient color) to determine whether behavioral performance on the fixation task depended on the type of ignored peripheral stimulus presented on each trial. All behavioral analyses only used fixation target trials that occurred during the stimulus presentation period, as this was the trial period of interest in neuroimaging analyses.

For all primary fMRI analyses, we focused on sets of retinotopically defined regions selected *a priori* based on previous reports establishing feature selectivity for color or motion (see above). Additionally, for completeness, we repeated these tests across each individual retinotopic ROI (results presented in supplemental information). For analyzing BOLD responses, we first compared univariate responses across stimulus conditions by computing separate two-way ANOVAs for each feature-selective ROI, with stimulus feature (color/motion) and salience presence (present/absent) as factors (Fig. S2). Next, to determine the spatial selectivity of reconstructed spatial maps based on fMRI activation patterns, we computed a three-way repeated measures ANOVA to determine the spatial selectivity of neural modulations with location activation (salient location, opposite location, and aligned position in salience-absent conditions), ROI, and feature (motion/color) as factors (Fig. 3D). To directly test whether feature-selective ROIs represent salient locations more strongly when salience is defined by their preferred feature value, we computed a paired-samples T-Test on the difference between map activation on color-salience and motion-salience trials between color-selective and motion-selective ROIs (Fig. 4). In supplemental analyses, we compared the same difference across all individual ROIs using a one-way repeated-measures ANOVA with ROI as factor (Fig. 4B; Fig. S4B). Finally, we assessed the spatial selectivity of feature-selective responses by computing a two-way ANOVA with ROI and location (salient location, location opposite to salient stimulus, and aligned position in salience absent condition) as factors (Fig. 4C; Fig. S4C).

For our shuffling procedure, we used a random number generator that was seeded with a single value for all analyses. The seed number was randomly selected using an online random number generator (https://numbergenerator.org/random-8-digit-number-generator). Within each participant, averaged data within each condition were shuffled across conditions for each participant individually, and once shuffled, the statistical test of interest was recomputed over 1000 iterations. *P* values were derived by computing the percentage of shuffled test statistics that were greater than or equal to the measured test statistic. We controlled for multiple comparisons using the false discovery rate (Benjamini & Yekutieli, 2001) across all comparisons within an analysis when necessary. Error bars are standard error, unless noted otherwise.

### Data & Code Availability

All data supporting the conclusions of this report and all associated analysis scripts will be available on Open Science Framework and GitHub upon publication. To protect participant privacy, and in accordance with IRB-approved procedures, freely available data is limited to extracted timeseries for each voxel of each ROI for all scans of the study. Whole-brain ‘raw’ data will be made available from the authors upon reasonable request.

## Results

### Behavior

Participants continuously monitored a central fixation cross for brief changes in line segment length while viewing stimuli in the periphery (Fig. 2). Stimuli could either be full-screen arrays of colored static or grayscale moving dots, or flickering checkerboards. When stimuli were dot arrays, they typically contained a salient region (colored dots: a disc appeared in a different color, 180° away in HSV colorspace; moving dots: a disc contained dots moving in the opposing motion direction). This design requires attention to be maintained at fixation, and allows for bottom-up salience to be isolated and evaluated in response to the ignored, task-irrelevant peripheral stimulus as a function of the salience-defining feature. Across runs we adjusted fixation task difficulty to keep behavioral performance above chance and below ceiling and to maximize participant engagement (average response accuracy across conditions: 81.07% % ± 2.13%, mean ± SEM across participants; average miss rate across conditions: 26.49% ± 9.12%). Importantly, we observed no difference in response accuracy (*p* = 0.7; one-way permuted repeated-measures ANOVA) or miss rate (*p* = 0.29; one-way permuted repeated-measures ANOVA) as a function of peripheral stimulus type. Thus, any differences observed in multivariate activation patterns between conditions cannot be driven by differences in behavioral performance.

### Multivariate Spatial Representations

Next, we used a spatial inverted encoding model (IEM) to reconstruct spatial maps based on measured activation patterns from each ROI on each trial (see Figs. S1/2 for results from univariate analyses). We used data from an independent ‘mapping’ task (see methods; Sprague et al., 2018) to estimate a spatial encoding model for each voxel parameterized as a set of weights on smooth, overlapping spatial channels. Then, we inverted the set of encoding models across all voxels in each cluster of regions to reconstruct spatial maps based on activation profiles from the feature-salience task (Fig. 3A). This procedure generates a reconstructed image for each timepoint, which we then averaged within condition and across timepoints corresponding to 5 to 8 s after stimulus onset. The resulting images are well-established to show strong activation at locations corresponding to visual stimulation (e.g., where a checkerboard was presented; Sprague et al., 2019; Sprague, Itthipuripat, et al., 2018; Sprague & Serences, 2013). Indeed, when the ‘checkerboard’ trials were used for stimulus reconstruction, we observed strong representations of the salient stimulus location in all ROIs (Fig. 3B; Fig. S3A).

Having validated our method, we asked: does activation within these reconstructions additionally track locations made salient based on local differences in feature values? In our study, the entire visual field is equivalently stimulated (e.g., equal amount of motion energy or density of colored dots), so any nonuniform activation must be due to salience-related activation patterns within each ROI. Indeed, model-based reconstructions were able to track the salient location throughout the visual field on both motion- and color-salient trials (Fig. 3C). When the salient location was defined by dots moving in a different motion direction, the reconstructed spatial map from TO1/TO2 showed a stronger representation of the salient location than when the salient location was defined by static dots presented in a different color, and the converse result is apparent when examining spatial maps reconstructed from hV4/VO1/VO2 (Fig. 3C). Critically, in these trials, the feature values at each location are updated at 3 Hz, minimizing the possibility that reconstructions of salient locations emerge from a serendipitous selection of a specific local feature value.

To quantify the condition-specific stimulus representation within model-based reconstructions for each ROI, we computed the mean activation at the known position of the salient stimulus (see Sprague et al, 2018; Fig. 3B). For ROIs which compute spatial maps of salient location(s), we predict that reconstructions will show an enhanced representation of the salient stimulus position when compared to non-salient locations. Our design allows for two important comparisons to establish whether these salience computations occur. First, we can directly compare the activation in reconstructed spatial maps at the salient location to activation in the location on the opposite side of the fixation point (which contains non-salient ‘background’ dots with an equal amount of color/motion energy as the salient location). This comparison allows us to demonstrate that spatial maps highlight salient locations within each salient stimulus condition. Second, we can compare the mean activation of the salient location on salience-present trials to a randomly selected location of the reconstructed spatial map on salience-absent trials. This allows us to see if the map activation at the salient location was greater than map activation at an equivalent spatial location when viewing a uniform dot array with no salient position(s).

Map activation values were strongest at the salient location when a salient stimulus was presented, and weaker at non-salient locations (both when a salient stimulus was presented elsewhere, and when no salient stimulus was present at all; Fig. 3D). We compared map activation values across conditions, map locations, and ROIs using a three-way repeated-measures permuted ANOVA with stimulus feature (motion/color), activation location (salience-absent; salience-present: salient location; salience-present: opposite location), and ROI (TO1/TO2; hV4/VO1/VO2) as factors. This analysis indicated that there was a main effect of activation location (*p* < 0.001), a two-way interaction between stimulus feature and ROI (*p* = 0.007), and a three-way interaction between all three factors (*p* = 0.001). All other comparisons were non-significant (*p* > 0.05).

Within hV4/VO1/VO2, we observed a significant difference between map activation at the salient location and opposite location on color salience-present trials (*p* < 0.001, permuted paired samples T-test; Fig. 3D) as well as a significant difference between the color-defined salient location and map activation on salience-absent colored dot trials (*p* = 0.008, permuted paired samples T-test). Map activation at the salient location was significantly greater than zero (*p* < 0.001; permuted one-sample T-test).

We found complementary results in TO1/TO2 when using data from the motion stimulus conditions (Fig. 3D). We observed a significant difference in map activation between the salient location defined by motion and the opposite location on salience-present trials (*p* < 0.001, permuted paired samples T-test) in addition to a significant difference between the map activation at the salient location and map activation from similar locations on trials when no salient location was defined (*p* < 0.001, permuted paired samples T-test). Map activation at salient locations defined by motion was also greater than zero (*p* < 0.001; permuted one-sample T-test). Altogether, these results suggest that activation patterns in these regions reflect the image-computable salience of the corresponding location in the visual field.

### Comparing salience computations across feature dimensions

Thus far, we have shown that color- and motion-selective regions each compute a representation of the location of a salient stimulus defined by feature contrast. If these regions act as neural dimension maps which each individually compute representations of salient locations defined by their preferred feature value, we expect to observe a more efficient extraction of salient locations when the salience-defining feature matches the region’s preferred feature value. We tested this by computing a pixelwise difference between reconstructed spatial maps from each ROI when salience was defined based on color and when salience was defined based on motion (Fig. 4A), along with the same difference between the salience-absent control conditions. Zero values (white) indicate that the map activation is equal between stimulus features, while positive (red)/negative (blue) values near the salient location indicate that the map preferentially extracts salient locations when the salience defining feature is motion or color, respectively. If each ROI selectively identifies salient locations based on its preferred feature value, then these difference maps will show greater (absolute) differences at the salient location than other locations. However, if instead feature selectivity and salience computations each independently and additively impact spatial maps, then these difference maps should show no spatial structure (particularly, no difference between salient and non-salient locations).

Qualitatively comparing these difference maps (Fig. 4A), it is apparent that both ROIs show a stronger difference at the salient location than the opposite location, consistent with a local and specialized computation of salient locations based on each region’s preferred feature values. The localized differences at the salient location are in opposite directions (TO1/TO2: positive/red; hV4/VO1/VO2: negative/blue), as expected if motion (color)-selective ROIs more efficiently extract salient locations defined by motion (color) feature contrast than color (motion) feature contrast.

We quantified the degree to which each ROI selectively computes salience based on its preferred feature value by extracting activation values from these difference maps at the salient location, the opposite location, and the ‘aligned’ location in the salience-absent condition and computing map activation difference scores based on the regions’ feature preferences (TO1/TO2: motion – color; hV4/VO1/VO2: color – motion; Fig. 4B). A two-way repeated measures ANOVA with location (salient location, opposite location, salience-absent) and ROI as factors revealed a significant main effect of location (*p* = 0.002). Follow-up comparisons in TO1/TO2 demonstrate that map activation was greater at the salient location than the opposite location (*p* = 0.016, permuted paired samples T-test) and the salience-absent location (*p* < 0.001, permuted paired samples T-test) and was the only location with activation greater than zero (*p* = 0.005, permuted one-sample T-test). In hV4/VO1/VO2, the same tests did not reveal significant differences between the salient location and the opposite location (*p* = 0.074, permuted paired samples T-test) or the salient-absent position (*p* = 0.556, permuted paired samples T-test). However, only the salient location had map activation differences greater than zero after FDR corrections (*p* = 0.001, permuted paired samples T-test). Together, these results suggest that each ROI selectively indexes a salient location when salience is defined based on its preferred feature value.

Finally, to directly compare salience computations between regions, we computed a salience modulation index (SMI). This was formalized as the difference between map activation at the salient location between the motion and color conditions, where positive values indicate a stronger response to the motion-defined salient location, negative values indicate a stronger response to the color-defined salient location, and zero means no difference between conditions. SMI reliably differed between motion- and color-selective ROIs (Fig. 4C; *p* < 0.001; permuted paired-samples T-test). This indicates that feature-selective ROIs preferentially compute salience based on their preferred feature dimension, and further supports the proposal that these retinotopically-defined regions act as neural dimension maps within priority map theory.

## Discussion

In the present study, our goal was to determine whether visual cortex computes spatial maps representing salient locations based on specific feature dimensions within feature-selective retinotopic regions (Fig. 1)—a key, yet untested, prediction of priority map theory. We probed this question by reconstructing spatial maps based on fMRI activation patterns measured while participants viewed, but ignored, stimuli containing salient locations based on different feature dimensions (Fig. 2). Our results show that salient location representations in color-selective regions hV4/VO1/VO2 and motion-selective regions TO1/TO2 are modulated by bottom-up feature salience even when top-down attention is kept at fixation (Fig. 3). Representations were selectively enhanced when the salience-defining feature matched the preferred feature of a given region and these enhancements occurred at the *salient location* (Fig. 4). These results provide strong evidence that these retinotopic cortical regions act as ‘neural feature dimension maps’, confirming an important untested prediction of priority map theory.

In previous studies in humans and nonhuman primates, neural correlates of salience and/or priority maps have been identified in several regions, including: LGN (Kastner et al., 2006; Poltoratski et al., 2017), V1 (Li, 2002; Poltoratski et al., 2017; Wang et al., 2022; Zhang et al., 2012), extrastriate visual cortex (including V4/hV4; Adam & Serences, 2021; Bogler et al., 2011, 2013; Burrows & Moore, 2009; Mazer & Gallant, 2003; Poltoratski et al., 2017; Sprague, Itthipuripat, et al., 2018; Sprague & Serences, 2013), LIP/IPS (Adam & Serences, 2021; Bisley & Goldberg, 2003, 2006; Chen et al., 2020; Gottlieb et al., 1998; Jerde et al., 2012; Sprague, Itthipuripat, et al., 2018), FEF (Bichot & Schall, 1999; Schall et al., 1995; Schall & Hanes, 1993), substantia nigra (Basso & Wurtz, 2002), pulvinar (Shipp, 2003), and SC (Basso & Wurtz, 1998; Fecteau & Munoz, 2006; White et al., 2017). Across these studies, activity in neurons/voxels tuned for salient and/or relevant locations is greater than activity in neurons/voxels tuned towards nonsalient and/or nonrelevant locations. However, many of these previous studies are limited by focusing on a single salience-defining feature and/or by relying on sparse single-unit recordings from one or a handful of cells within one or a small number of brain regions in non-human primates. Such studies necessarily face difficulty comparing across different brain regions and assessing activation profiles across the entire region (and thus, the entire visual field). Here, we overcame these limitations by implementing a multivariate IEM which allowed us to reconstruct activation profiles across a map of the entire visual field from each timepoint’s measured activation pattern in each ROI (Fig. 3A). Additionally, by manipulating the salience-defining stimulus feature dimension across trials (Fig. 2) while simultaneously measuring fMRI activation patterns across multiple feature-selective ROIs (Fig. S1/2), we established that the region best representing a salient location depends on the salience-defining feature (Figs. 3-4).

As mentioned above, there is extensive, and often conflicting, evidence for priority maps implemented in different structures throughout the brain. With the seemingly redundant computations of priority across regions, how is information across maps ultimately leveraged to guide attention? We expect measurements of feature dimension maps like those identified in the current study can be used to disentangle these various accounts by establishing which regions act to integrate information about salient locations based on combinations of features. One testable prediction of the priority map framework is that activation profiles in a feature-agnostic priority map should reflect some computation over the activation profiles measured across individual feature dimension maps, such as: linear combination (Wolfe, 1984), winner-take-all processes (Itti & Koch, 2001), and probabilistic integration (Eckstein, 2017). While such a test is not possible in our current study, future work incorporating stimuli with multiple salient locations with different degrees of stimulus-defined salience may be able to better disentangle the roles of various putative priority maps in guiding visual attention based on stimulus properties.

Aspects of priority map theory are invoked by foundational models of behavioral performance on visual search tasks (Duncan & Humphreys, 1989; Folk et al., 1992; Müller et al., 1995; Theeuwes, 2010). For example, when a subject is tasked with searching for one shape among a homogeneous array of other shapes (e.g., a circle among squares), search performance is slower if one of the distracting stimuli is presented in a different color (Theeuwes, 1992), and slowing is greater for larger target/distractor color discrepancies (Duncan & Humphreys, 1989; Theeuwes, 1992; Wolfe & Horowitz, 2017). Influential cognitive models posit that the color distractor results in greater activation in a priority map, which slows search for the shape-defined target stimulus. Indeed, reconstructed spatial representations of target and distractor stimuli measured from extrastriate visual and parietal cortex during an adapted version of this task show enhanced neural representations for color-defined distracting stimuli (Adam & Serences, 2021). While cognitive priority map models have offered parsimonious explanations of changes in discrimination performance, RTs, and gaze trajectories, they often disagree on when and how salient items capture attention (Luck et al., 2021). One central issue limiting the ability to adjudicate among competing models is that there is no well-established method for quantitatively measuring now neural activity indexes the relative salience of different aspects of stimulus displays (Chang et al., 2021; Gaspelin & Luck, 2021; Pearson et al., 2021). Our findings may offer a practical solution to this challenge. Here, we demonstrate that neural representations of feature-based salience can simultaneously be tracked across multiple feature-selective regions, and previous work has identified similar salience representations using stimuli varying in luminance contrast across multiple retinotopic regions (Sprague et al., 2018). Together, this work establishes a framework for empirically estimating neural representations of visual salience across cortical processing stages, including feature dimension maps. Using these methods, future work can investigate how competing stimuli made salient by different feature dimensions are represented within retinotopic maps and how the relative strength of those representations – and how they interact with one another within and between maps – may explain aspects of how salient stimuli capture attention measured using behavioral methods.

Our results show that feature-selective retinotopic ROIs compute a stronger representation of the salient location defined based on their preferred feature dimension as compared to a non-preferred feature dimension. However, each ROI still represents the salient location, even when made salient by the non-preferred feature value (Fig. 3). We speculate that this is due to feedback from higher-order regions (e.g., parietal or frontal cortex) that aggregate salience maps across individual feature dimensions to guide attention to important locations in the scene. Because the observers’ task required careful fixation and the stimulus was always irrelevant, such automatic extraction of salient locations was never used by the participant to guide covert or overt attention. However, it may be the case that the automatic identification of salient scene locations results in automatic feedback signals across retinotopic cortex, similar to widespread retinotopic effects of cued spatial attention observed previously (e.g., Gandhi et al., 1999; Itthipuripat et al., 2019; Sprague, Itthipuripat, et al., 2018; Sprague & Serences, 2013; Tootell et al., 1998). Indeed, reconstructions based on parietal cortex activation patterns show representations of the salient location, as do all other retinotopic regions studied (Fig. S3). Importantly, only feature-selective regions TO1/TO2 and hV4/VO1/VO2 show a systematic change in the representation of the salient location as a function of the salience-defining feature, supporting their role as neural feature dimension maps despite their weaker representation of a salient location based on a non-preferred feature.

While this study provides evidence that specialized computations support the identification of salient locations based on different feature values, there remain some important limitations to this work that will be addressed in future studies. First, in this study, we required participants direct attention to the fixation point to perform a very challenging visual discrimination task near their performance threshold. This design choice aimed to minimize the role of peripheral covert attention in driving activation within the measured ROIs, but it also eliminated the opportunity to relate our measured neural activation profiles with behavioral performance. Future studies employing visual search displays (e.g., Adam & Serences, 2021) will better disentangle the impact of feature-selective salience computations on performance of demanding cognitive tasks (e.g., those requiring visual search in the presence of salient distractors). Additionally, to maximize our ability to detect representations of salient locations, we used stimuli of fixed size but random location defined by 100% feature contrast (color: opposite hues in HSV colorspace; motion: opposite motion directions). Future studies which parametrically manipulate the size, number, and feature contrast of salient stimulus locations in similar stimulus displays (e.g., Bogler et al, 2013; Zhang et al, 2012; Burrows & Moore, 2009) could enable both comparison of reconstructed spatial maps across various levels of stimulus salience and in-depth forward modeling of salience-related computations and their associated nonlinearities based on local feature contrast input to each voxel’s receptive field (e.g., Hughes et al., 2019; Kay, Winawer, Rokem, et al., 2013; Yildirim et al., 2018). Finally, future studies are required to test whether activation profiles in these neural feature dimension maps are equivalently sensitive to screen regions made salient by increases and decreases in feature intensity (e.g., motion speed, color saturation), which is a manipulation that has previously been effectively used to dissociate location-specific activation driven by stimulus intensity and local salience computations (Betz et al, 2013).

In summary, we found that feature-selective retinotopic ROIs compute maps of stimulus salience primarily based on feature contrast within their preferred feature dimension, confirming a key untested prediction of priority map theory. These results identify feature-selective retinotopic regions as the neural correlates of feature dimension maps within the priority map framework and support a new approach for probing the neural computations supporting visual cognition.

## Acknowledgments

This work was supported by an Alfred P Sloan Research Fellowship, an Nvidia Hardware Grant, a UC Santa Barbara Academic Senate Research Grant, and US Army Research Office Cooperative Agreement W911NF-19-2-0026 for the Institute for Collaborative Biotechnologies. We thank Kirsten Adam and John Serences for helpful comments on a draft of the manuscript.

## Supplemental Information

**Figure S1.**
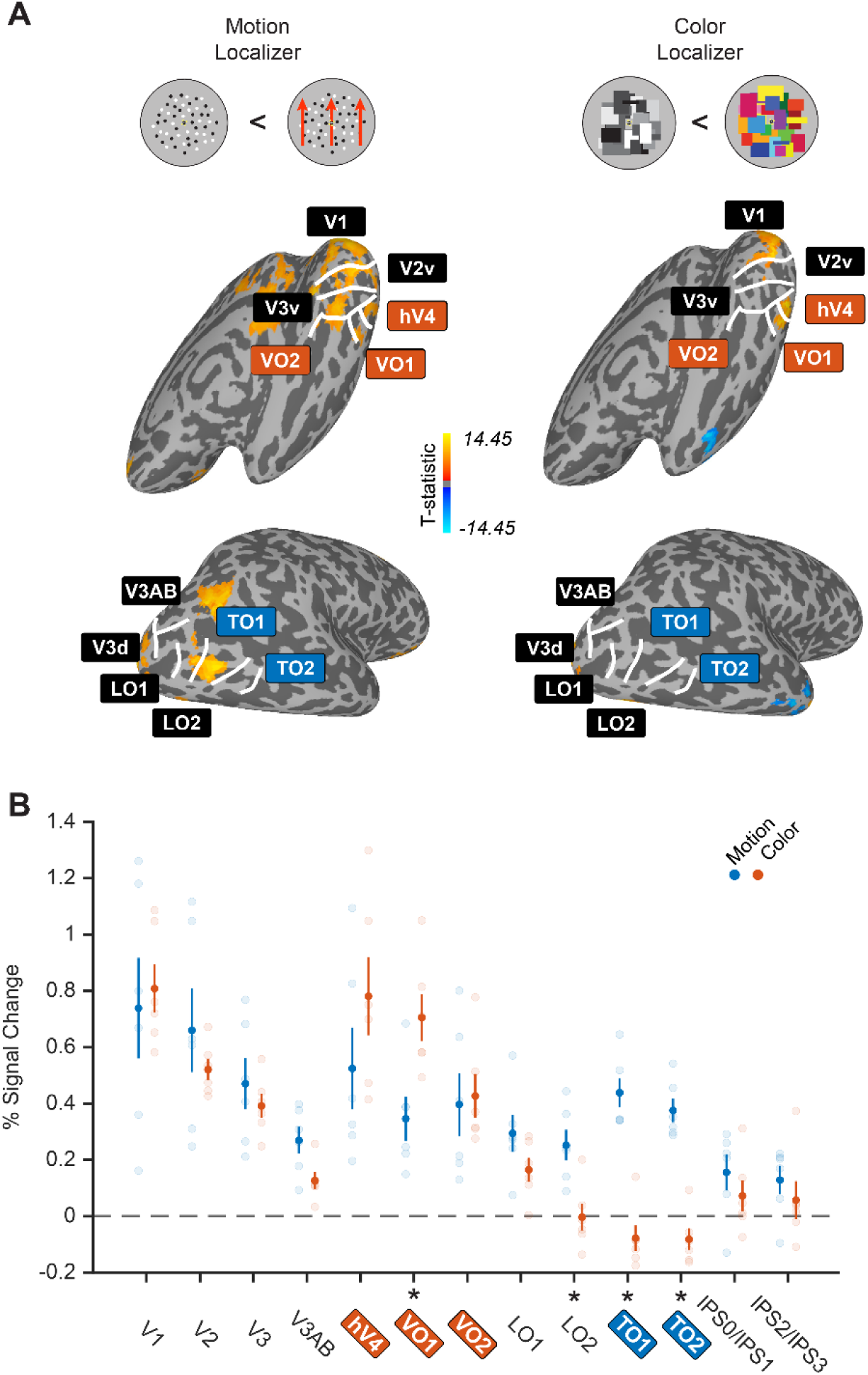
Localizer-evoked activation in retinotopically-defined ROIs confirms feature selectivity. **A.** We scanned N = 6 of our participants on a feature localizer task which required careful attention to the fixation point while moving/static grayscale dots (motion) or colored/grayscale rectangles (color) stimuli appeared in blocks of 12 s, separated by 12 s ITIs. Example stimulus displays for each condition in the motion/color localizers. Plotted t-statistic maps from an example subject (sub011) for comparisons between moving/static and color/grayscale stimulus displays overlayed with retinotopic ROIs drawn based on polar angle reversals identified with our retinotopic mapping task (see Fig. S2 and Methods). Even though motion localizer contrasts show various significant activations throughout ventral & lateral retinotopic ROIs, there was a high degree of overlap with retinotopic region TO1. Color localizer activations strongly overlap with retinotopic regions hV4 and VO1, with minimal activation in lateral retinotopic regions, including TO1/TO2. Activation map based on T values from the colored vs grayscale (or motion vs static) contrast (*T* ≥ 6, cluster threshold of 40 neighboring voxels). **B.** We fit a GLM and extracted beta weights corresponding to each condition and plotted the contrasts (difference values) within each feature dimension (Color: colored – grayscale rectangles; Motion: moving – static dots) for each ROI. Permuted 2-way repeated measures ANOVA (ROI; localizer feature) showed a main effect of ROI (*p* = 0.013) but not localizer feature (*p* = 0.263). However, there was a significant interaction (*p* = 0.002). Notably, only regions with previously established feature-selectivity (color: VO1; motion: TO1, TO2) showed a significant difference between localizer types, with the exception of LO2, which also showed a preference for motion. However, this region shares a border with TO1, and so slight overlap between regions and/or imperfections in ROI definition may have led to an enhanced response to the motion localizer in LO2. While color- and motion-contrasts in hV4 and VO2 did not significantly differ from one another, qualitatively, hV4 was more sensitive to the color manipulation than the motion manipulation. See Figs. S2-S4 for results from individual ROIs, which show a qualitatively consistent result using only VO1 as computed using hV4, VO1, and VO2 as a combined color-selective ROI. * Indicates significant difference between motion and color contrasts at *p* < 0.05 after FDR correction. Datapoints and error bars show mean ± SEM across participants. See Table S1 for all statistical tests.

**Figure S2.**
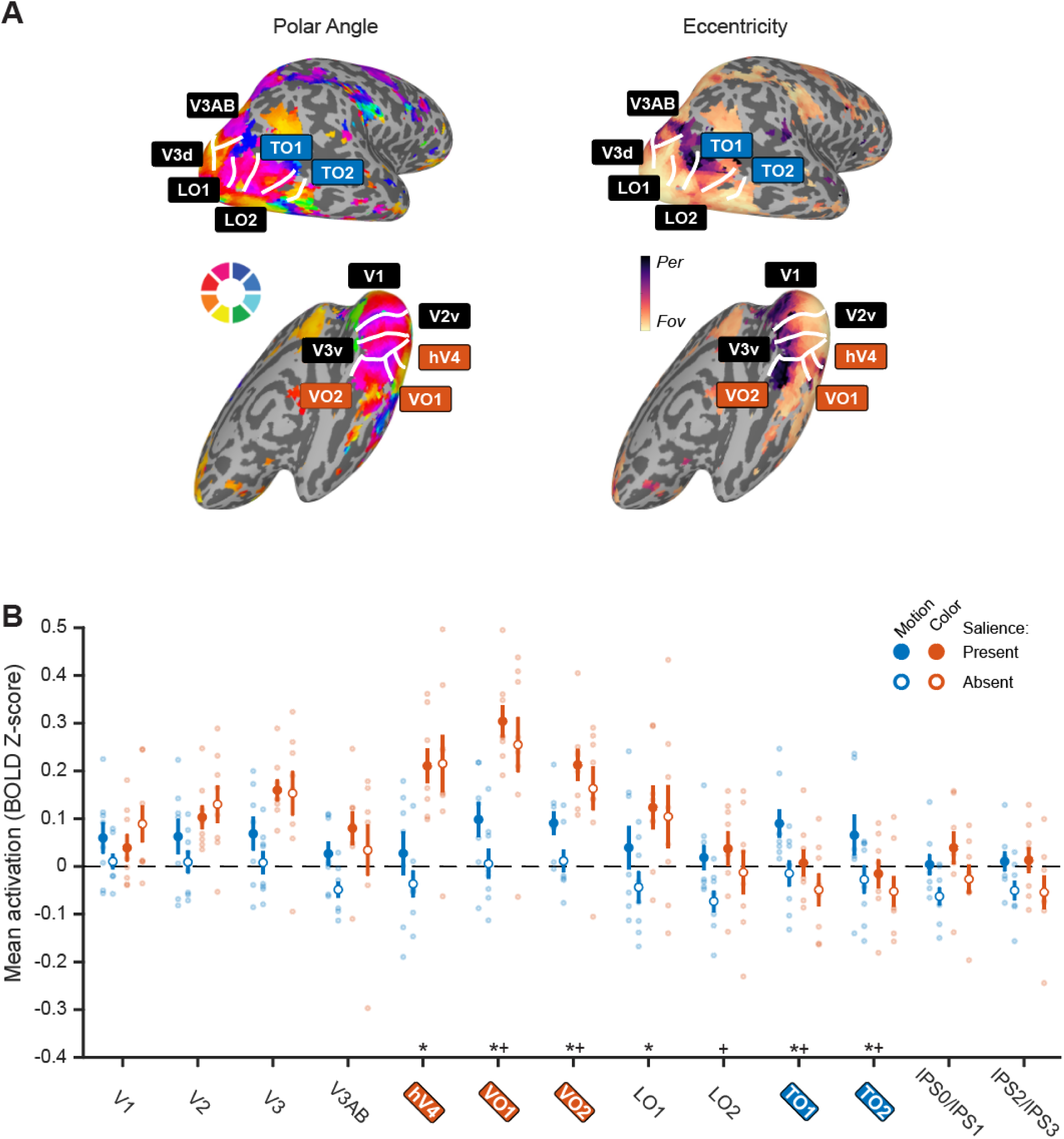
Retinotopic ROIs show univariate selectivity for stimulus feature. **A.** We identified ROIs based on polar angle and eccentricity maps measured using a voxel receptive field-based retinotopic mapping procedure (see Methods). Based on previous reports, we defined hV4/VO1/VO2 as color-selective and TO1/TO2 as motion-selective regions (Amano et al., 2009; Brewer et al., 2005; Huk et al., 2002; Wade et al., 2002). We confirmed these ROIs aligned with color- and motion-selective patches defined using traditional functional localizer approaches (Fig. S1A). Polar angle (left) and eccentricity (right) maps shown for right hemisphere of example participant (sub011. Fov: foveal, Per: peripheral). **B.** Average activation for retinotopic ROIs (5-8 s after stimulus onset) was greatest when the dot array was defined based on the preferred feature of that region for individual feature-selective ROIs. Additionally, trials containing a salient dot patch resulted in greater mean univariate activation. Permuted 3-way repeated measures ANOVA (ROI; salience; feature) showed a main effect of ROI (*p* < 0.001), salience (*p* = 0.024), and feature (*p* = 0.013). There were also significant interactions between salience and ROI (*p* < 0.001) as well as feature and ROI (*p* < 0.001). All other interactions did not reach significance (*p* > 0.1). All feature-selective ROIs showed a main effect of feature, supporting the stimulus preferences of each region. * indicates main effect of feature (color/motion), + indicates main effect of salience presence (salient location present/absent), and × indicates interaction (permuted 2-way repeated-measures ANOVA). Error bars reflect SEM across participants. See Table S2 for all statistical tests.

**Figure S3.**
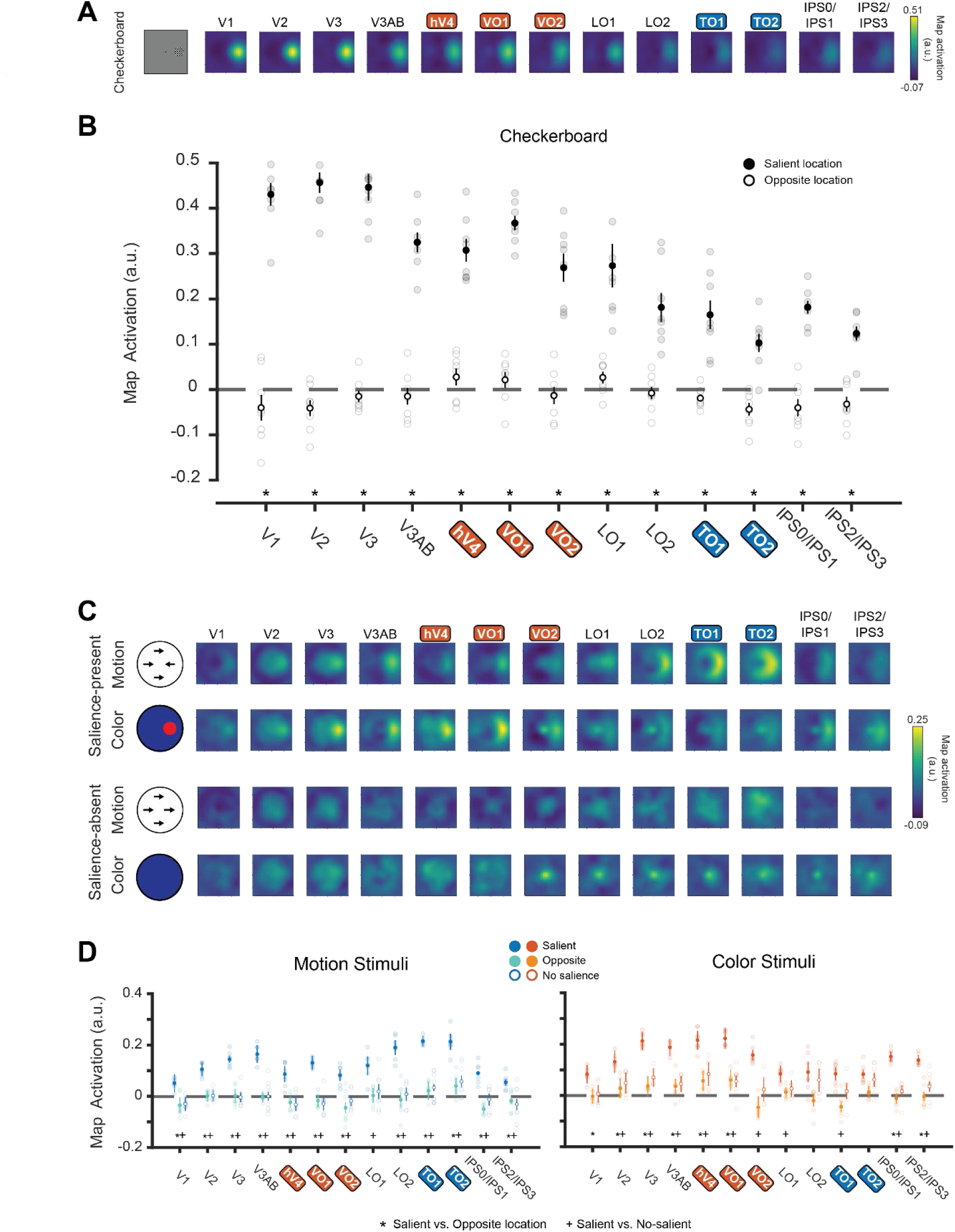
Reconstructed spatial maps track salient stimulus location. **A:** Reconstructions of checkerboard stimuli from each individual retinotopic ROI. Qualitatively, there was a strong response to the checkerboard stimulus across all ROIs. **B.** We quantified reconstructions by computing the mean map activation at the aligned checkerboard stimulus location and at the location on the opposite side of fixation within each ROI. Permuted 2-way repeated measures ANOVA (ROI and location) showed a significant main effect of ROI (*p* < 0.001), location (*p* < 0.001) and interaction (*p* < 0.001). Map activation was stronger at the location of the checkerboard as compared to the opposite location in all ROIs. * indicates significant difference based on permuted paired T-test, corrected for multiple comparisons with FDR. **C.** Reconstructions were computed for the salience-present and -absent conditions for both features (color/motion; as in Fig. 3). Qualitatively, the salient location was highlighted in feature-selective ROIs when the salient location was defined by motion, but not by color, with the converse result observed in color-selective ROIs. On salience-absent trials, no location is reliably highlighted in average reconstructions. **D**: Using data from each stimulus condition, we identified whether enhanced reconstruction responses were localized to the salient location by comparing mean salient location activation to the mean activation of the position opposite of the salient location, as well as the mean activation of the ‘aligned’ position of the salience-absent condition. On trials with motion-defined stimuli, activation in motion-selective ROIs was greatest at the location of the salient motion patch. When stimuli were defined by static colorful dots, activation in the color-selective ROI was greatest at the location of the salient color stimulus. * indicates significant difference between salient and opposite locations, and + indicates significant difference between the salient location and salience-absent reconstructions. Both set of tests are based on permuted paired T-test and corrected for multiple comparisons with FDR. 3-way permuted repeated-measures ANOVA (ROI; location; stimulus feature) identified a significant main effect of location (*p* < 0.001), 2-way interaction between feature and ROI (*p* < 0.001), and a 3-way interaction (*p* < 0.001). In all panels, error bars are SE across participants (n = 8). For all statistical comparisons, see Table S3.

**Figure S4.**
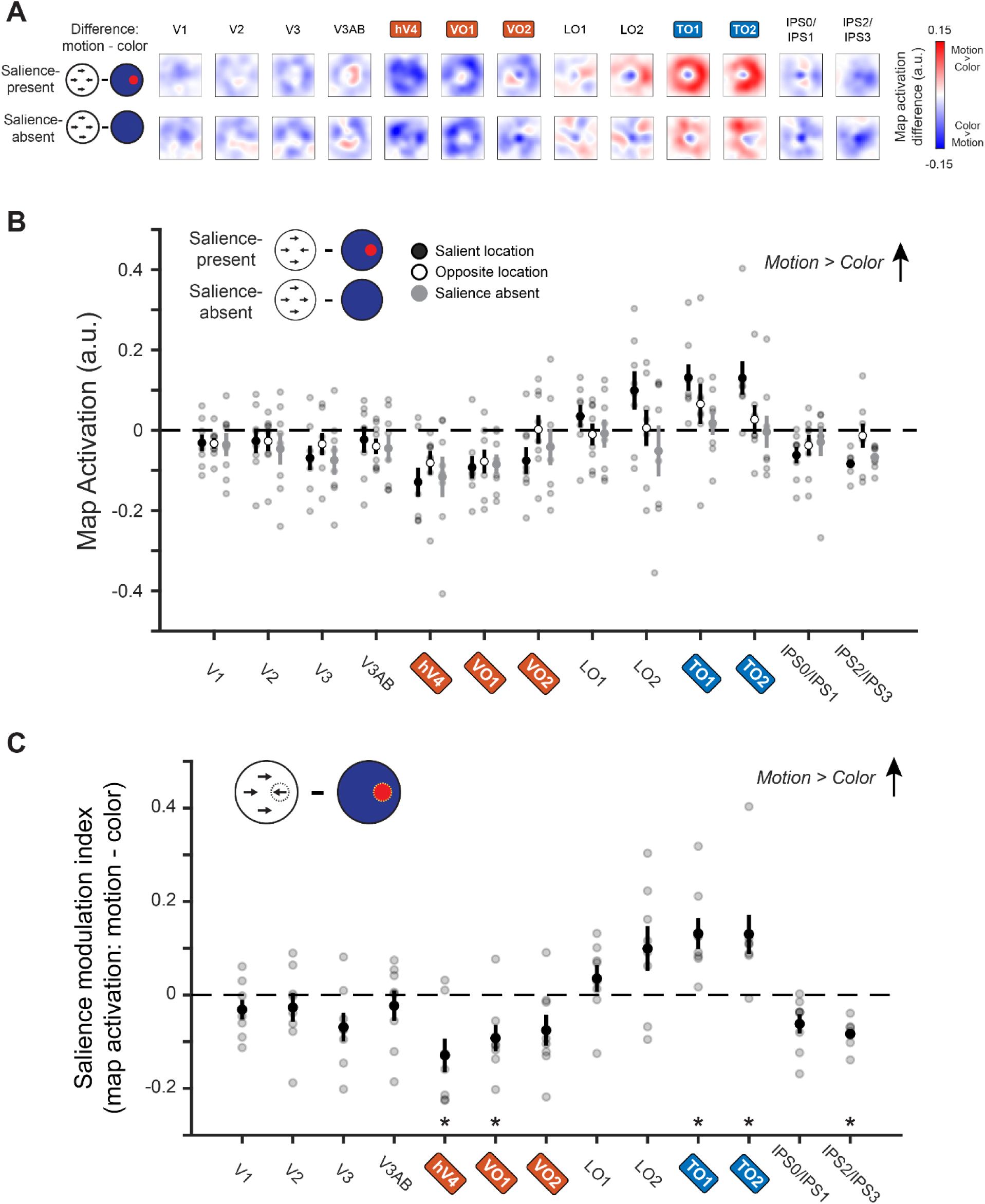
Neural dimension maps selectively index salience based on their preferred feature. **A:** We directly compared reconstructed spatial maps for each ROI between trials where the salient location was defined by motion to those where the salient location was defined by color by computing the pixelwise difference (motion – color). For comparison, we also computed these maps for salience-absent trials. Positive values indicate a region more strongly represents a salient location based on motion over color, and negative values indicate the opposite. Data presented as in Fig. 4A. **B:** To compare feature selectivity across spatial locations and salience-presence conditions, we extracted values of each ROI’s difference map at the salient location, opposite location, and the salience-absent condition. Absolute difference map activation was greater at the salient location than the opposite location on salience-present trials than at a random location on salience-absent trials, and this effect depended on ROI (2-way permuted repeated-measures ANOVA with factors of location and ROI; significant main effect of ROI *p* < 0.001 and interaction *p* < 0.001). **C**: Difference map activation (A) computed at the salient location was only reliably different from 0 in feature-selective ROIs, such that motion-selective regions indexed salience more strongly when it was defined by motion than by color, and vice versa for color-selective regions. Asterisk indicates significant difference based on permuted one-sample T-test, *p* < 0.05. * indicates significance difference from zero. Error bars reflect SEM across participants. See Table S4 for all statistical tests.

**Table S1.** Localizer statistical tests (related to Figure S1). P-values for all localizer comparisons and each ROI. Comparisons include one-sample tests for each feature contrast (motion and color) and between feature dimensions. Bold numbers indicate significant differences after FDR correction for all comparisons (*q* = 0.05). Italicized numbers indicate significance before FDR corrections using *α* = 0.05.

| Figure S1 | Motion Contrast | Color Contrast | Motion vs Color |
| --- | --- | --- | --- |
| V1 | **0.009** | **< 0.001** | 0.762 |
| V2 | **0.007** | **< 0.001** | 0.402 |
| V3 | **0.004** | **< 0.001** | 0.556 |
| V3ab | **0.002** | **0.008** | 0.088 |
| hV4 | **0.015** | **0.002** | 0.190 |
| VO1 | **0.007** | **< 0.001** | *0.022* |
| VO2 | **0.016** | **0.003** | 0.870 |
| LO1 | **0.006** | **0.012** | 0.104 |
| LO2 | **0.006** | 0.946 | *0.030* |
| TO1 | **< 0.001** | 0.149 | **< 0.001** |
| TO2 | **< 0.001** | 0.088 | **< 0.001** |
| hV4/VO1/VO2 | **0.011** | **< 0.001** | 0.246 |
| TO1/TO2 | **< 0.001** | 0.154 | **0.004** |
| IPS0/IPS1 | 0.060 | 0.251 | 0.410 |
| IPS2/IPS3 | 0.052 | 0.434 | 0.478 |

**Table S2.** Univariate statistical tests (related to Figure 2 and Figure S2). P-values for all univariate comparisons and each ROI. Comparisons include one-sample tests for each feature/salience combination as well as comparisons between feature dimension for the salience-present and -absent conditions. Bold numbers indicate significant differences after FDR correction for all comparisons (*q* = 0.05). Italicized numbers indicate significance before FDR corrections using *α* = 0.05.

| Figure 2/S2 | Motion |  |  | Color |  |  | Motion and Color |  |
| --- | --- | --- | --- | --- | --- | --- | --- | --- |
|  | Salient | Salience absent | Salient vs salience absent | Salient | Salience absent | Salient vs Salience absent | Color vs motion (salience) | Color vs motion (salience absent) |
| V1 | 0.116 | 0.551 | 0.128 | 0.224 | 0.055 | 0.086 | 0.676 | 0.108 |
| V2 | 0.136 | 0.716 | 0.064 | **0.005** | *0.013* | 0.322 | 0.404 | *0.034* |
| V3 | 0.095 | 0.749 | 0.054 | **< 0.001** | *0.014* | 0.902 | 0.056 | *0.018* |
| V3ab | 0.326 | *0.024* | *0.030* | 0.058 | 0.547 | 0.468 | 0.154 | 0.248 |
| hV4 | 0.569 | 0.240 | 0.072 | **0.001** | *0.009* | 0.936 | **0.002** | *0.006* |
| VO1 | *0.032* | 0.861 | *0.008* | **< 0.001** | **0.003** | 0.310 | **0.002** | **0.002** |
| VO2 | *0.008* | 0.642 | *0.006* | **< 0.001** | *0.009* | 0.230 | **0.002** | *0.016* |
| LO1 | 0.417 | 0.242 | *0.028* | *0.031* | 0.159 | 0.708 | 0.064 | *0.016* |
| LO2 | 0.500 | *0.014* | *0.02* | 0.339 | 0.799 | 0.240 | 0.680 | 0.156 |
| TO1 | *0.019* | 0.611 | *0.012* | 0.803 | 0.199 | 0.180 | *0.016* | 0.340 |
| TO2 | 0.171 | 0.390 | *0.022* | 0.635 | 0.149 | 0.354 | *0.018* | 0.442 |
| TO1/TO2 | *0.038* | 0.509 | **0.012** | 0.964 | 0.170 | 0.174 | **0.016** | 0.312 |
| hV4/VO1 /VO2 | 0.085 | 0.771 | **0.024** | **< 0.001** | **0.005** | 0.538 | **0.002** | **0.004** |
| IPS0/1 | 0.868 | *0.014* | *0.046* | 0.289 | 0.417 | 0.196 | 0.280 | 0.484 |
| IPS2/3 | 0.635 | *0.044* | 0.062 | 0.637 | 0.169 | 0.180 | 0.884 | 0.982 |

**Table S3.** Statistical tests on multivariate reconstructed spatial maps (related to Figure 3 and Figure S3). P-values for comparisons at specific locations in reconstructed spatial maps for each ROI. Comparisons include one-sample T-tests for each motion, color, and checkerboard stimulus at the location of the salient stimulus, the opposite location (loc), and the salience-absent condition. Two-sample T-tests were conducted for all condition combinations (salience loc vs opposite loc, salience loc vs salience absent, opposite loc vs salience absent). Bold numbers indicate significant differences after FDR correction for all comparisons (*q* = 0.05). 0 indicates comparisons where *p* < 0.001. Italicized numbers indicate significance before FDR corrections using *α* = 0.05.

| Figure 3/S3 | Motion |  |  |  |  |  | Color |  |  |  |  |  | Checkerboard |  |  |
| --- | --- | --- | --- | --- | --- | --- | --- | --- | --- | --- | --- | --- | --- | --- | --- |
|  | Salient Location | Opposite Location | Saliency-absent | Salient loc vs Saliency-absent loc vs no sing | Salient loc vs opposite loc location | Saliency-absent vs opposite loc | Salient Location | Opposite location | Saliency-absent | Salient loc vs Saliency-absent | Salient loc vs opposite loc | Saliency-absent vs opposite loc | Checker location | Opposite location | Checker loc vs opposite loc |
| V1 | 0.195 | 0.290 | 0.309 | **0.002** | **0** | 0.768 | 0.057 | 0.940 | 0.869 | **0** | 0.004 | 0.656 | **0** | 0.200 | **0** |
| V2 | **0.005** | 0.966 | 0.899 | **0** | **0.002** | 0.928 | *0.018* | 0.493 | 0.277 | **0** | **0** | 0.268 | **0** | 0.052 | **0** |
| V3 | **0** | 0.941 | 0.828 | **0** | **0** | 0.674 | **0** | 0.391 | 0.093 | **0** | **0** | **0.002** | **0** | 0.278 | **0** |
| V3ab | **0.002** | 0.828 | 0.999 | **0** | **0** | 0.842 | **0** | 0.274 | 0.186 | **0** | **0** | 0.718 | **0** | 0.443 | **0** |
| hV4 | *0.028* | 0.283 | 0.235 | 0.008 | **0.008** | 0.678 | **0** | 0.195 | 0.107 | **0** | **0** | 0.392 | **0** | 0.180 | **0** |
| VO1 | **0.002** | 0.485 | 0.305 | **0** | **0** | 0.672 | **0** | 0.198 | 0.043 | **0** | **0** | 0.944 | **0** | 0.259 | **0** |
| VO2 | *0.010* | 0.071 | 0.413 | **0.006** | **0** | 0.084 | **0** | 0.313 | 0.628 | **0.01** | **0** | 0.022 | **0** | 0.512 | **0** |
| LO1 | *0.014* | 0.941 | 0.546 | *0.040* | **0.004** | 0.422 | *0.034* | 0.556 | 0.437 | **0.030** | **0** | 0.458 | **0.001** | 0.079 | **0.002** |
| LO2 | **0** | 0.652 | 0.747 | **0.002** | **0** | 0.542 | 0.056 | 0.407 | 0.227 | 0.284 | 0.004 | 0.068 | **0.001** | 0.588 | **0.004** |
| TO1 | **0** | 0.575 | *0.038* | **0** | **0** | 0.738 | *0.013* | 0.082 | 0.614 | **0.012** | **0** | 0.076 | **0.001** | *0.046* | **0** |
| TO2 | **0** | 0.253 | *0.040* | **0** | **0** | 0.532 | **0.002** | 0.592 | 0.055 | **0.262** | 0.066 | 0.206 | **0.001** | **0.017** | **0** |
| TO1/TO2 | **0** | 0.325 | **0.004** | **0** | **0** | 0.408 | **0.004** | 0.237 | 0.164 | **0.006** | **0.002** | 0.054 | **0** | **0.009** | **0** |
| hV4/VO1<br>/VO2 | **0.001** | 0.194 | 0.151 | **0** | **0** | 0.796 | **0** | 0.345 | 0.176 | **0.008** | **0** | 0.546 | **0** | *0.024* | **0** |
| IPS0/1 | **0** | *0.028* | 0.617 | **0.004** | **0** | 0.210 | **0** | 0.575 | 0.486 | **0** | **0** | 0.146 | **0** | 0.075 | **0** |
| IPS2/3 | **0.003** | 0.215 | 0.202 | **0.004** | **0.002** | 0.678 | **0** | 0.853 | 0.114 | **0** | **0** | 0.076 | **0** | 0.087 | **0.002** |

**Table S4.** Feature-selective map activation statistical tests (related to Figure 4 and Figure S4). P-values for comparisons at specific locations within feature-difference spatial maps for each ROI. Comparisons include one-sample T-tests at the location of the salient stimulus, the opposite location, and the salience-absent maps. Two-sample T-tests were conducted for all location combinations (salience vs opposite, salience vs absent, opposite vs absent). Bold numbers indicate significant differences after FDR correction for all comparisons (*q* = 0.05). Italicized numbers indicate significance before FDR corrections using *α* = 0.05.

| Figure 4B-C/S4 | Salient Location | Opposite Location | Salience absent | Salient loc vs opposite loc | Salient loc vs salience-absent | Opposite loc vs salience-absent |
| --- | --- | --- | --- | --- | --- | --- |
| V1 | 0.176 | *0.041* | 0.252 | 0.938 | 0.788 | 0.894 |
| V2 | 0.415 | 0.318 | 0.282 | 0.948 | 0.472 | 0.556 |
| V3 | 0.061 | 0.246 | 0.079 | 0.244 | 0.866 | *0.046* |
| V3ab | 0.496 | 0.081 | 0.219 | 0.480 | 0.590 | 0.83 |
| hV4 | *0.009* | *0.029* | 0.055 | 0.178 | 0.820 | 0.322 |
| VO1 | *0.014* | *0.034* | *0.010* | 0.508 | 0.736 | 0.548 |
| VO2 | 0.057 | 0.954 | 0.385 | 0.096 | 0.554 | 0.176 |
| LO1 | 0.253 | 0.713 | 0.824 | 0.062 | 0.208 | 0.944 |
| LO2 | 0.077 | 0.902 | 0.444 | **0.004** | *0.016* | 0.316 |
| TO1 | *0.005* | 0.233 | 0.576 | 0.106 | **0.006** | 0.350 |
| TO2 | *0.017* | 0.476 | 0.927 | **0.002** | **0.002** | 0.260 |
| TO1/TO2 | **0.005** | 0.180 | 0.529 | **0.016** | **< 0.001** | 0.174 |
| hV4/VO1<br>/VO2 | **0.001** | *0.049* | *0.038* | 0.074 | 0.556 | 0.388 |
| IPS0/1 | *0.020* | 0.194 | 0.434 | 0.318 | 0.188 | 0.81 |
| IPS2/3 | **< 0.001** | 0.667 | **< 0.001** | *0.030* | 0.354 | 0.106 |

## References

1. Adam, K. C. S., & Serences, J. T. (2021). History Modulates Early Sensory Processing of Salient Distractors. Journal of Neuroscience, 41(38), 8007–8022. https://doi.org/10.1523/JNEUROSCI.3099-20.2021

2. Albright, T. D. (1993). Cortical processing of visual motion. Reviews of Oculomotor Research, 5, 177–201.

3. Amano, K., Wandell, B. A., & Dumoulin, S. O. (2009). Visual field maps, population receptive field sizes, and visual field coverage in the human MT+ complex. Journal of Neurophysiology, 102(5), 2704–2718.

4. Awh, E., Belopolsky, A. V., & Theeuwes, J. (2012). Top-down versus bottom-up attentional control: A failed theoretical dichotomy. Trends in Cognitive Sciences, 16(8), 437–443.

5. Bacon, W. F., & Egeth, H. E. (1994). Overriding stimulus-driven attentional capture. Perception & Psychophysics, 55(5), 485–496. https://doi.org/10.3758/BF03205306

6. Baker, D. H., Vilidaite, G., Lygo, F. A., Smith, A. K., Flack, T. R., Gouws, A. D., & Andrews, T. J. (2021). Power contours: Optimising sample size and precision in experimental psychology and human neuroscience. Psychological Methods, 26, 295–314. https://doi.org/10.1037/met0000337

7. Bartels, A., & Zeki, S. (2000). The architecture of the colour centre in the human visual brain: New results and a review. European Journal of Neuroscience, 12(1), 172–193.

8. Basso, M. A., & Wurtz, R. H. (1998). Modulation of Neuronal Activity in Superior Colliculus by Changes in Target Probability. Journal of Neuroscience, 18(18), 7519–7534. https://doi.org/10.1523/JNEUROSCI.18-18-07519.1998

9. Basso, M. A., & Wurtz, R. H. (2002). Neuronal Activity in Substantia Nigra Pars Reticulata during Target Selection. Journal of Neuroscience, 22(5), 1883–1894. https://doi.org/10.1523/JNEUROSCI.22-05-01883.2002

10. Beck, D. M., & Kastner, S. (2005). Stimulus context modulates competition in human extrastriate cortex. Nature Neuroscience, 8(8), Article 8. https://doi.org/10.1038/nn1501

11. Benjamini, Y., & Yekutieli, D. (2001). The control of the false discovery rate in multiple testing under dependency. The Annals of Statistics, 29(4), 1165–1188. https://doi.org/10.1214/aos/1013699998

12. Bichot, N. P., & Schall, J. D. (1999). Effects of similarity and history on neural mechanisms of visual selection. Nature Neuroscience, 2(6), Article 6. https://doi.org/10.1038/9205

13. Bisley, J. W., & Goldberg, M. E. (2003). Neuronal activity in the lateral intraparietal area and spatial attention. Science, 299(5603), 81–86.

14. Bisley, J. W., & Goldberg, M. E. (2006). Neural Correlates of Attention and Distractibility in the Lateral Intraparietal Area. Journal of Neurophysiology, 95(3), 1696–1717. https://doi.org/10.1152/jn.00848.2005

15. Bisley, J. W., & Goldberg, M. E. (2010). Attention, Intention, and Priority in the Parietal Lobe. Annual Review of Neuroscience, 33(1), 1–21. https://doi.org/10.1146/annurev-neuro-060909-152823

16. Bogler, C., Bode, S., & Haynes, J.-D. (2011). Decoding Successive Computational Stages of Saliency Processing. Current Biology, 21(19), 1667–1671. https://doi.org/10.1016/j.cub.2011.08.039

17. Bogler, C., Bode, S., & Haynes, J.-D. (2013). Orientation pop-out processing in human visual cortex. NeuroImage, 81, 73–80. https://doi.org/10.1016/j.neuroimage.2013.05.040

18. Brainard, D. H. (1997). The Psychophysics Toolbox. Spatial Vision, 10(4), 433–436. https://doi.org/10.1163/156856897X00357

19. Brewer, A. A., Liu, J., Wade, A. R., & Wandell, B. A. (2005). Visual field maps and stimulus selectivity in human ventral occipital cortex. Nature Neuroscience, 8(8), 1102–1109.

20. Burrows, B. E., & Moore, T. (2009). Influence and Limitations of Popout in the Selection of Salient Visual Stimuli by Area V4 Neurons. Journal of Neuroscience, 29(48), 15169–15177. https://doi.org/10.1523/JNEUROSCI.3710-09.2009

21. Carrasco, M. (2011). Visual attention: The past 25 years. Vision Research, 51(13), 1484–1525. https://doi.org/10.1016/j.visres.2011.04.012

22. Chang, S., Niebur, E., & Egeth, H. E. (2021). Standing out in a small crowd: The role of display size in attracting attention. Visual Cognition, 29(9), 587–591. https://doi.org/10.1080/13506285.2021.1918810

23. Chen, X., Zirnsak, M., Vega, G. M., Govil, E., Lomber, S. G., & Moore, T. (2020). Parietal Cortex Regulates Visual Salience and Salience-Driven Behavior. Neuron.

24. Conway, B. R., Moeller, S., & Tsao, D. Y. (2007). Specialized Color Modules in Macaque Extrastriate Cortex. Neuron, 56(3), 560–573. https://doi.org/10.1016/j.neuron.2007.10.008

25. Cook, E. P., & Maunsell, J. H. R. (2002). Attentional Modulation of Behavioral Performance and Neuronal Responses in Middle Temporal and Ventral Intraparietal Areas of Macaque Monkey. Journal of Neuroscience, 22(5), 1994–2004. https://doi.org/10.1523/JNEUROSCI.22-05-01994.2002

26. Dumoulin, S. O., & Wandell, B. A. (2008). Population receptive field estimates in human visual cortex. Neuroimage, 39(2), 647–660.

27. Duncan, J., & Humphreys, G. W. (1989). Visual search and stimulus similarity. Psychological Review, 96(3), 433–458. https://doi.org/10.1037/0033-295X.96.3.433

28. Eckstein, M. P. (2011). Visual search: A retrospective. Journal of Vision, 11(5), 14–14. https://doi.org/10.1167/11.5.14

29. Eckstein, M. P. (2017). Probabilistic Computations for Attention, Eye Movements, and Search. Annual Review of Vision Science, 3(1), 319–342. https://doi.org/10.1146/annurev-vision-102016-061220

30. Fecteau, J. H., & Munoz, D. P. (2006). Salience, relevance, and firing: A priority map for target selection. Trends in Cognitive Sciences, 10(8), 382–390. https://doi.org/10.1016/j.tics.2006.06.011

31. Gandhi, S. P., Heeger, D. J., & Boynton, G. M. (1999). Spatial attention affects brain activity in human primary visual cortex. Proceedings of the National Academy of Sciences, 96(6), 3314–3319. https://doi.org/10.1073/pnas.96.6.3314

32. Gaspelin, N., & Luck, S. J. (2021). Progress and remaining issues: A response to the commentaries on Luck et al. (2021). Visual Cognition, 0(0), 1–7. https://doi.org/10.1080/13506285.2021.1979705

33. Gottlieb, J. P., Kusunoki, M., & Goldberg, M. E. (1998). The representation of visual salience in monkey parietal cortex. Nature, 391(6666), Article 6666. https://doi.org/10.1038/35135

34. Hallenbeck, G. E., Sprague, T. C., Rahmati, M., Sreenivasan, K. K., & Curtis, C. E. (2021). Working memory representations in visual cortex mediate distraction effects. Nature Communications, 12(1), Article 1. https://doi.org/10.1038/s41467-021-24973-1

35. Huang, L., & Pashler, H. (2007). A Boolean map theory of visual attention. Psychological Review, 114, 599– 631. https://doi.org/10.1037/0033-295X.114.3.599

36. Hughes, A. E., Greenwood, J. A., Finlayson, N. J., & Schwarzkopf, D. S. (2019). Population receptive field estimates for motion-defined stimuli. NeuroImage, 199, 245–260. https://doi.org/10.1016/j.neuroimage.2019.05.068

37. Huk, A. C., Dougherty, R. F., & Heeger, D. J. (2002). Retinotopy and functional subdivision of human areas MT and MST. Journal of Neuroscience, 22(16), 7195–7205.

38. Itthipuripat, S., Sprague, T. C., & Serences, J. T. (2019). Functional MRI and EEG index complementary attentional modulations. Journal of Neuroscience, 39(31), 6162–6179.

39. Itti, L., & Koch, C. (2000). A saliency-based search mechanism for overt and covert shifts of visual attention. Vision Research, 40(10), 1489–1506. https://doi.org/10.1016/S0042-6989(99)00163-7

40. Itti, L., & Koch, C. (2001). Computational modelling of visual attention. Nature Reviews Neuroscience, 2(3), 194.

41. Jerde, T. A., Merriam, E. P., Riggall, A. C., Hedges, J. H., & Curtis, C. E. (2012). Prioritized Maps of Space in Human Frontoparietal Cortex. The Journal of Neuroscience, 32(48), 17382–17390. https://doi.org/10.1523/JNEUROSCI.3810-12.2012

42. Kastner, S., Schneider, K. A., & Wunderlich, K. (2006). Beyond a relay nucleus: Neuroimaging views on the human LGN. Progress in Brain Research, 155, 125–143.

43. Katsuki, F., & Constantinidis, C. (2014). Bottom-Up and Top-Down Attention: Different Processes and Overlapping Neural Systems. The Neuroscientist, 20(5), 509–521. https://doi.org/10.1177/1073858413514136

44. Kay, K. N., Winawer, J., Mezer, A., & Wandell, B. A. (2013). Compressive spatial summation in human visual cortex. Journal of Neurophysiology, 110(2), 481–494.

45. Kay, K. N., Winawer, J., Rokem, A., Mezer, A., & Wandell, B. A. (2013). A Two-Stage Cascade Model of BOLD Responses in Human Visual Cortex. PLOS Computational Biology, 9(5), e1003079. https://doi.org/10.1371/journal.pcbi.1003079

46. Li, Z. (2002). A saliency map in primary visual cortex. Trends in Cognitive Sciences, 6(1), 9–16. https://doi.org/10.1016/S1364-6613(00)01817-9

47. Mackey, W. E., Winawer, J., & Curtis, C. E. (2017). Visual field map clusters in human frontoparietal cortex. Elife, 6, e22974.

48. Martínez-Trujillo, J. C., & Treue, S. (2002). Attentional Modulation Strength in Cortical Area MT Depends on Stimulus Contrast. Neuron, 35(2), 365–370. https://doi.org/10.1016/S0896-6273(02)00778-X

49. Mazer, J. A., & Gallant, J. L. (2003). Goal-related activity in V4 during free viewing visual search. Evidence for a ventral stream visual salience map. Neuron, 40(6), 1241–1250. https://doi.org/10.1016/s0896-6273(03)00764-5

50. Moran, J., & Desimone, R. (1985). Selective Attention Gates Visual Processing in the Extrastriate Cortex. Science, 229(4715), 782–784. https://doi.org/10.1126/science.4023713

51. Mulckhuyse, M., Belopolsky, A. V., Heslenfeld, D., Talsma, D., & Theeuwes, J. (2011). Distribution of Attention Modulates Salience Signals in Early Visual Cortex. PLOS ONE, 6(5), e20379. https://doi.org/10.1371/journal.pone.0020379

52. Mullen, K. T. (2019). The response to colour in the human visual cortex: The fMRI approach. Current Opinion in Behavioral Sciences, 30, 141–148. https://doi.org/10.1016/j.cobeha.2019.08.001

53. Müller, H. J., Heller, D., & Ziegler, J. (1995). Visual search for singleton feature targets within and across feature dimensions. Perception & Psychophysics, 57(1), 1–17. https://doi.org/10.3758/BF03211845

54. O’Connell, T. P., & Chun, M. M. (2018). Predicting eye movement patterns from fMRI responses to natural scenes. Nature Communications, 9(1), 5159. https://doi.org/10.1038/s41467-018-07471-9

55. Ogawa, T., & Komatsu, H. (2004). Target Selection in Area V4 during a Multidimensional Visual Search Task. Journal of Neuroscience, 24(28), 6371–6382. https://doi.org/10.1523/JNEUROSCI.0569-04.2004

56. Ogawa, T., & Komatsu, H. (2006). Neuronal dynamics of bottom-up and top-down processes in area V4 of macaque monkeys performing a visual search. Experimental Brain Research, 173(1), 1–13. https://doi.org/10.1007/s00221-006-0362-5

57. Pearson, D., Watson, P., & Le Pelley, M. E. (2021). How do competing influences of selection history interact? A commentary on Luck et al. (2021). *Visual Cognition*, 0(0), 1–4. https://doi.org/10.1080/13506285.2021.1912234

58. Pelli, D. G. (1997). The VideoToolbox software for visual psychophysics: Transforming numbers into movies. Spatial Vision, 10(4), 437–442. https://doi.org/10.1163/156856897X00366

59. Poltoratski, S., Ling, S., McCormack, D., & Tong, F. (2017). Characterizing the effects of feature salience and top-down attention in the early visual system. Journal of Neurophysiology, 118(1), 564–573. https://doi.org/10.1152/jn.00924.2016

60. Poltoratski, S., & Tong, F. (2020). Resolving the Spatial Profile of Figure Enhancement in Human V1 through Population Receptive Field Modeling. Journal of Neuroscience, 40(16), 3292–3303. https://doi.org/10.1523/JNEUROSCI.2377-19.2020

61. Reynolds, J. H., & Desimone, R. (2003). Interacting Roles of Attention and Visual Salience in V4. Neuron, 37(5), 853–863. https://doi.org/10.1016/S0896-6273(03)00097-7

62. Schall, J. D., & Hanes, D. P. (1993). Neural basis of saccade target selection in frontal eye field during visual search. Nature, 366(6454), Article 6454. https://doi.org/10.1038/366467a0

63. Schall, J. D., Morel, A., King, D. J., & Bullier, J. (1995). Topography of visual cortex connections with frontal eye field in macaque: Convergence and segregation of processing streams. Journal of Neuroscience, 15(6), 4464–4487. https://doi.org/10.1523/JNEUROSCI.15-06-04464.1995

64. Scotti, P. S., Chen, J., & Golomb, J. D. (2021). An enhanced inverted encoding model for neural reconstructions (p. 2021.05.22.445245). https://doi.org/10.1101/2021.05.22.445245

65. Serences, J. T., & Yantis, S. (2006). Selective visual attention and perceptual coherence. Trends in Cognitive Sciences, 10(1), 38–45.

66. Shipp, S. (2003). The functional logic of cortico–pulvinar connections. Philosophical Transactions of the Royal Society of London. Series B: Biological Sciences, 358(1438), 1605–1624. https://doi.org/10.1098/rstb.2002.1213

67. Sprague, T. C., Adam, K. C., Foster, J. J., Rahmati, M., Sutterer, D. W., & Vo, V. A. (2018). Inverted encoding models assay population-level stimulus representations, not single-unit neural tuning. ENeuro, 5(3).

68. Sprague, T. C., Boynton, G. M., & Serences, J. T. (2019). Inverted encoding models estimate sensible channel responses for sensible models. https://doi.org/10.1101/642710

69. Sprague, T. C., Ester, E. F., & Serences, J. T. (2014). Reconstructions of information in visual spatial working memory degrade with memory load. Current Biology, 24(18), 2174–2180.

70. Sprague, T. C., Ester, E. F., & Serences, J. T. (2016). Restoring latent visual working memory representations in human cortex. Neuron, 91(3), 694–707.

71. Sprague, T. C., Itthipuripat, S., Vo, V. A., & Serences, J. T. (2018). Dissociable signatures of visual salience and behavioral relevance across attentional priority maps in human cortex. Journal of Neurophysiology, 119(6), 2153–2165.

72. Sprague, T. C., & Serences, J. T. (2013). Attention modulates spatial priority maps in the human occipital, parietal and frontal cortices. Nature Neuroscience, 16(12), 1879.

73. Swisher, J. D., Halko, M. A., Merabet, L. B., McMains, S. A., & Somers, D. C. (2007). Visual Topography of Human Intraparietal Sulcus. The Journal of Neuroscience, 27(20), 5326–5337. https://doi.org/10.1523/JNEUROSCI.0991-07.2007

74. Theeuwes, J. (1992). Perceptual selectivity for color and form. Perception & Psychophysics, 51(6), 599– 606. https://doi.org/10.3758/BF03211656

75. Tootell, R. B. H., Hadjikhani, N., Hall, E. K., Marrett, S., Vanduffel, W., Vaughan, J. T., & Dale, A. M. (1998). The Retinotopy of Visual Spatial Attention. Neuron, 21(6), 1409–1422. https://doi.org/10.1016/S0896-6273(00)80659-5

76. Treisman, A. (1998). The perception of features and objects. In Visual attention (pp. 26–54). Oxford University Press.

77. Treisman, A. M., & Gelade, G. (1980). A feature-integration theory of attention. Cognitive Psychology, 12(1), 97–136.

78. Wandell, B. A., Dumoulin, S. O., & Brewer, A. A. (2007). Visual field maps in human cortex. Neuron, 56(2), 366–383.

79. Wang, L., Huang, L., Li, M., Wang, X., Wang, S., Lin, Y., & Zhang, X. (2022). An awareness-dependent mapping of saliency in the human visual system. NeuroImage, 247, 118864. https://doi.org/10.1016/j.neuroimage.2021.118864

80. White, B. J., Berg, D. J., Kan, J. Y., Marino, R. A., Itti, L., & Munoz, D. P. (2017). Superior colliculus neurons encode a visual saliency map during free viewing of natural dynamic video. Nature Communications, 8(1), Article 1. https://doi.org/10.1038/ncomms14263

81. Winawer, J., & Witthoft, N. (2015). Human V4 and ventral occipital retinotopic maps. Visual Neuroscience, 32, E020. https://doi.org/10.1017/S0952523815000176

82. Wolfe, J. M. (1994). Guided search 2.0 a revised model of visual search. Psychonomic Bulletin & Review, 1(2), 202–238.

83. Wolfe, J. M., & Horowitz, T. S. (2004). What attributes guide the deployment of visual attention and how do they do it? Nature Reviews Neuroscience, 5(6), 495.

84. Wolfe, J. M., & Horowitz, T. S. (2017). Five factors that guide attention in visual search. Nature Human Behaviour, 1(3), 1–8. https://doi.org/10.1038/s41562-017-0058

85. Yildirim, F., Carvalho, J., & Cornelissen, F. W. (2018). A second-order orientation-contrast stimulus for population-receptive-field-based retinotopic mapping. NeuroImage, 164, 183–193. https://doi.org/10.1016/j.neuroimage.2017.06.073

86. Yu, X., Zhou, Z., Becker, S. I., Boettcher, S. E. P., & Geng, J. J. (2023). Good-enough attentional guidance. Trends in Cognitive Sciences, 27(4), 391–403. https://doi.org/10.1016/j.tics.2023.01.007

87. Zhang, X., Zhaoping, L., Zhou, T., & Fang, F. (2012). Neural Activities in V1 Create a Bottom-Up Saliency Map. Neuron, 73(1), 183–192. https://doi.org/10.1016/j.neuron.2011.10.035

